# Design and execution of a Verification, Validation, and Uncertainty Quantification plan for a numerical model of left ventricular flow after LVAD implantation

**DOI:** 10.1101/2021.11.11.468169

**Authors:** Alfonso Santiago, Constantine Butakoff, Beatriz Eguzkitza, Richard A. Gray, Karen May-Newman, Pras Pathmanathan, Vi Vu, Mariano Vázquez

## Abstract

**Background:** left ventricular assist devices (LVADs) are implantable pumps that act as a life support therapy for patients with severe heart failure. Despite improving the survival rate, LVAD therapy can carry major complications. Particularly, the flow distortion introduced by the LVAD in the left ventricle (LV) may induce thrombus formation. While previous works have used numerical models to study the impact of multiple variables in the intra-LV stagnation regions, a comprehensive validation analysis has never been executed. The main goal of this work is to present a model of the LV-LVAD system and to design and follow a verification, validation and uncertainty quantification (VVUQ) plan based on the ASME V&V40 and V&V20 standards to ensure credible predictions.

**Methods:** The experiment used to validate the simulation is the SDSU cardiac simulator, a bench mock-up of the cardiovascular system that allows mimicking multiple operation conditions for the heart-LVAD system. The numerical model is based on Alya, the BSC’s in-house platform for numerical modelling. Alya solves the Navier-Stokes equation with an Arbitrarian Lagrangian-Eulerian (ALE) formulation in a deformable ventricle and includes pressure-driven valves, a 0D Windkessel model for the arterial output and a LVAD boundary condition modeled through a dynamic pressure-flow performance curve. The designed VVUQ plan involves: *(a)* a risk analysis and the associated credibility goals; *(b)* a verification stage to ensure correctness in the numerical solution procedure; *(c)* a sensitivity analysis to quantify the impact of the inputs on the four quantities of interest (QoIs) (average aortic root flow 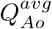, maximum aortic root flow 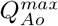, average LVAD flow 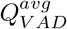, and maximum LVAD flow 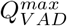); *(d)* an uncertainty quantification using six validation experiments that include extreme operating conditions.

**Results:** Numerical code verification tests ensured correctness of the solution procedure and numerical calculation verification showed small numerical errors. The total Sobol indices obtained during the sensitivity analysis demonstrated that the ejection fraction, the heart rate, and the pump performance curve coefficients are the most impactful inputs for the analysed QoIs.

The Minkowski norm is used as validation metric for the uncertainty quantification. It shows that the midpoint cases have more accurate results when compared to the extreme cases. The total computational cost of the simulations was above 100 [core-years] executed in around three weeks time span in Marenostrum IV supercomputer.

**Conclusions:** This work details a novel numerical model for the LV-LVAD system, that is supported by the design and execution of a VVUQ plan created following recognised international standards. We present a methodology demonstrating that stringent VVUQ according to ASME standards is feasible but computationally expensive.

## 1 Introduction

Over 5 million people suffer heart failure (HF) in the U.S. alone, with ~1 million new cases diagnosed annually [1]. Heart transplant is the recommended treatment for the 10% of these patients in Stage D [2] condition. Despite this, there are only 2000 organs yearly available for transplant [3], sufficient for only 0.4% of these patients. The limited organ availability is making left ventricular assist device (LVAD) therapy a leading treatment for the remaining 99.6% patients, with a ~90% chance of 1-year survival [4].

Mortality and HF status following LVAD implantation are primarily associated with inefficient unloading of the left ventricle and persistence of right ventricular dysfunction [5]. Optimization of LVAD speed is routinely performed for patients post-implant with a ramp study. Transthoracic echocardiography measurements of cardiac geometry and function are made while LVAD speed is slowly increased over a wide range [6]. The final pump speed is selected by balancing overall cardiac output, efficiency of left ventricle (LV) unloading, and preserving flow pulsatility [5]. Several variables are assessed from standard ultrasound views including LV end-diastolic dimension, LV end-systolic diameter, frequency of Aortic Valve (AoV) opening, degree of valve regurgitation, right ventricle (RV) systolic pressure, blood pressure, and heart rate (HR) at each speed setting. In addition, LVAD pump power, pulsatility index and flow are recorded. For the Thoratec HeartMate II, the ramp speed protocol starts at a speed of 8*k*[*rpm*] and increases by 400[*rpm*] increments every 2 minutes until a speed of 12*k*[*rpm*] is reached. As LVAD speed is increased, the LV volume decreases, as does the frequency of AoV opening and flow pulsatility. Excessive LV unloading at higher LVAD speeds increases the demand on the right heart, causing tricuspid regurgitation and also possibly produce suction events which disrupt the flow into the LVAD inflow cannula.

The clinical practice for LVAD speed selection first ensures that the hemodynamics are compatible with life, *e.g*. a mean arterial pressure greater than 65 mmHg [6] and a minimum cardiac index of 2.2[*L/min/m*2] of body surface area (BSA) [5] To optimize LV unloading, the interventricular septum position should not bow towards either the left or right. If these conditions are met, the LVAD speed is selected that achieves intermittent AoV opening while maintaining no more than mild mitral regurgitation or aortic insufficiency [7]. *De novo* AoV insufficiency development in LVAD patients is linked to lack of AoV opening. Interestingly, AoV insufficiency occurred in the majority of LVAD patients (66%) whose AoVs remained closed during support, but rarely (8%) in those whose AoVs opened regularly [6].

A patient specific LVAD speed calibration is important for ensuring appropriate cardiovascular support and minimizing the frequency of adverse events related to long-term support. However, the ramp echo study is not performed routinely after the first month post-implant, due to the unjustified expense and inconvenience. A computational tool that predicts cardiac output and aortic valve opening for the subject’s characteristics could reduce the ramp study requirements as well as contribute to speed adjustments required over a long-term, even supporting a speed adjustment paradigm that contributes to recovery. As LVADs are a life support therapy that carry a high risk for the patient, the Food and Drug Administration (FDA) rank them in the most exhaustive level of control (Class III) before approving their commercialisation. Therefore, using computational models to predict the behaviour of these devices or to guide design decisions should be accompanied with a stringent validation process to ensure credible results. Validating computational models is a whole challenge by itself, but recent published guidelines tackle this problem.

While scientific computing has undergone extraordinary increases in sophistication, a fundamental disconnect exists between simulations and practical applications. While most simulations are deterministic, engineering applications have many sources of uncertainty arising from a number of sources such as subject variability, initial conditions or system surroundings. Furthermore, the numerical model itself can introduce large uncertainties due to the assumptions and the numerical approximations employed [8]. Without forthrightly estimating the total uncertainty in a prediction, decision makers will be ill advised. To address this issue, multiple standards for industrial guidance has been published like the American Society of Mechanical Engineers (ASME) V&V codes [9, 10, 11].

Extensive modeling studies dealing with multiple LVAD factors like ventricular size [12], cannula implantation position [13], implantation depth [14, 15, 16] or angulation [17] exist but none of them provide credibility evidence as suggested in the recent ASME V&V40 [10], nor are any guided by AMSE V&V20 [11] which was developed over 10 years ago for demonstrating credibility of computational fluid dynamics (CFD) models. The reason for this is that such a validation requires a thorough comparison of the simulation results against bench or animal experiment measurements and hundreds of executions of the numerical model, which involves a large computational cost.

To our knowledge, this is the first paper describing such a comprehensive computational LVAD setup and using ASME verification, validation and uncertainty quantification (VVUQ) standards to assess the model credibility. While our final goal is to predict intra-LV stagnation biomarkers, this manuscript is focused on the credibility assessment of the numerical model. The contributions of this manuscript combines:

1. A deformable ventricle numerical model using a unidirectional fluid-structure interaction (FSI) and 0D aortic impedance model (as in [15]).
2. A novel pressure-driven valve model for mitral and aortic valves.
3. A dynamic pressure-flow (also called H-Q) performance curve for the LVAD boundary condition.
4. A VVUQ plan designed and executed following the ASME V&V40 and V&V20 standards [10, 11].
5. A set of validation metrics to quantify the differences between the simulation and the experiment and that describe the aortic valve flow and total cardiac output.

### Note on the nomenclature

The words “model” and “experiment” may work for any simpler representation of a more complex system (*e.g*. animal, bench or numerical model/experiment). For the sake of brevity, we refer to the bench model/experiment simply as “experiment” and to the numerical model/experiment as “simulation”, except if stated otherwise.

## 2 Methods

### 2.1 Description of the benchtop model

The experiments were performed with the San Diego State University (SDSU) cardiac simulator (CS), shown in Fig. 2. This CS is a mock circulation loop of the heart and the circulatory system with an apically implanted LVAD (Abbott HeartMate II) that has been reported previously in [18, 19]. It involves a transparent model of the dilated LV based on an idealised geometry, immersed in a water-filled tank and connected to an external circulatory loop mimicking the systemic circulation. The tank is fully watertight, so when the piston pump generates negative pressure, the LV expands to the end diastolic volume (EDV). The LV used is manufactured from platinum-cured silicone rubber (Young’s modulus *E* = 620[*kPa*] at 100% elongation and ultimate tensile strength of *P_max_* = 5.52[*MPa*] at 400% elongation). Porcine valves were used in both the aortic position (26[*mm*] Medtronic 305 Cinch) and the mitral position (25[*mm*] 6625 Carpentier Edwards Bioprosthesis). Tygon tubing (16[*mm*] diameter) replaced the HeartMate II outflow graft and was connected to the ascending aorta at a 90[°] angle approximately 15[*mm*] distal to the aortic root. The circulating fluid was a viscosity-matched blood analog consisting of 40% glycerol (viscosity of *μ* = 3.72[*cP*] at 20[°*C*]) and saline [20]. The systemic circulation can be represented with a 3-element Windkessel model with 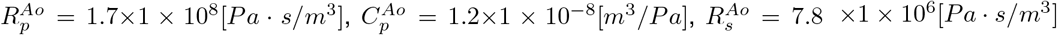, following the method in [21]. During diastole the silicone ventricle dilates allowing the filling with fluid from the atrial chamber. During systole the ventricle contracts expelling fluid through the aortic valve.

**Fig. 2:**
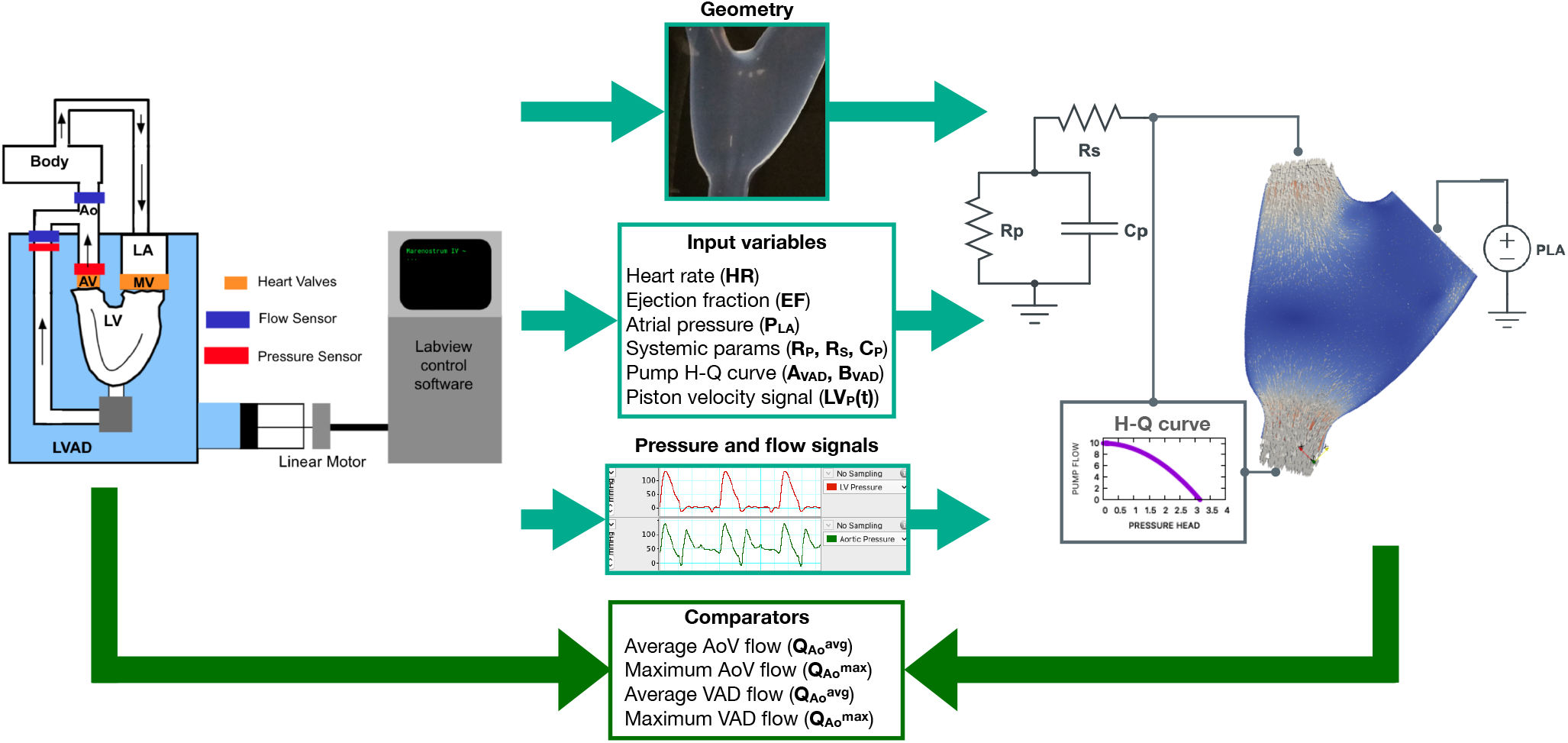
Leftmost side: Schematic of the experiment setup. LA: left atrium, MV: Mitral valve, AV: Aortic valve, Ao: aorta. Rightmost side: Schematic of the numerical model, including the lumped models for the boundary conditions. Middle: variables extracted and processed form the experiment and used in the simulation and comparators used for the uncertainty quantification (UQ).

Through this process the LVAD is continuously extracting fluid out of the LV. If the LVAD speed is high enough, the aortic valve remains hemodynamically closed during systole.

Flows are measured for the total aortic flow *Q_TAo_* (Transonic TS410 20PXL clamp-on ^1^) and the LVAD flow *Q_V AD_* (Transonic TS410 10PXL^2^). The aortic valve flow *Q_Ao_* is calculated as *Q_AoV_* = *Q_TAo_* – *Q_V AD_*. Pressure was also measured (Icumed TranspacIV ^3^) in the LV and the aortic root. Signals are amplified collected (ADinstruments Powerlab DAQ ^4^) and processed with LabChart (ADInstruments^5^) [22].

Two beating modes and three pump speeds are used for six validation experiments (Table 4). The beating mode 22[%]@68.42[*bpm*] has *EF* = 22[%] and *HR* = 68.42[*bpm*] with end systolic volume (ESV)=180[*cm*^3^] and EDV=230[*cm*^3^]. The beating mode 17[%]@61.18[*bpm*] has *EF* = 17[%] and *HR* = 61.18[*bpm*] with ESV=180[*cm*^3^] and EDV=216.86[*cm*^3^]. The Ejection Fraction (EF), the EDV, and the ESV are linked by the EF equation:

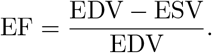

As the ESV is fixed by the silicone ventricle volume, there is a direct correlation between EF and EDV. From now on, we only characterise the case as a function of the EF. Both beating modes correspond to a New York heart association (NYHA) Class IV HF patient [23]. The pump speeds used for the validation points are 0[*rpm*], 8*k*[*rpm*] and 11*k*[*rpm*]. To avoid backflow, the LVAD outflow conduit were clamped in the 0[*rpm*] experiments so the system mimics a pre-LVAD baseline rather than a heart implanted with a LVAD turned off. The pressure-flow (also called H-Q) performance curves are experimentally retrieved for each pump speed (see Fig. 14) and used afterwards as input for the simulation.

### 2.2 Description of the numerical model

#### 2.2.1 Overall simulation pipeline

The computational domain is created from the exact same computer geometry used to manufacture the silicone ventricle (refer to Fig. 2). The unidirectional FSI requires the construction of two meshes. The solid mechanics mesh is created directly from the original geometry. The CFD mesh was created by closing the solid domain and extruding the inlets and outlets to ensure flow development. The meshes have 200*k* for the solid and 1.6*M* elements for the fluid. Special care was taken with the fluid mesh, including a boundary layer valid for *Re* < 4000[-]. The mesh is spatially discretised using linear tethrahedra, pyramids and pentagons. For the time discretisation, a first order trapezoidal rule with a time step of 0.00428[*s*] was used in every case. For a representation of the domain refer to Figs. 5 and 9.

To obtain a computationally inexpensive and accurate way of deforming the ventricle, a unidirectional FSI [24] approach is used to deform the LV (similarly as [15]). A pressure is imposed in the external solid domain which afterwards deform the CFD domain between the ESV and the EDV.

Once the simulation pipeline is completed, the input files are modified to work as a template. This template is used by Dakota server (DARE) (described in Section 2.2.6) for the sensitivity analysis (SA) and the UQ analysis.

#### 2.2.2 Description of the CFD solver

Here we briefly describe the numerical model used, highlighting only the novel components. The incompressible, Newtonian fluid is modelled by the Navier-Stokes equations in Alya, the Barcelona Supercomputing Center (BSC) in-house tool for simulations [25]. The Navier-Stokes equations are solved using an Arbitrarian Lagrangian-Eulerian (ALE) formulation, allowing the fluid domain to deform:

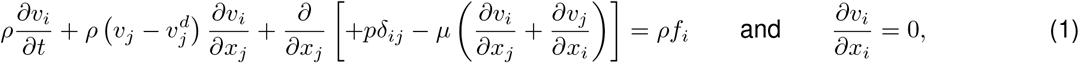

where *μ* is the dynamic viscosity of the fluid, *ρ* the density, *v_i_* the velocity, *p* is the mechanical pressure, *f_i_* the volumetric force term and 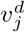 is the domain velocity. As the fluid domain deforms due to the imposed boundary displacements, the deformation for the inner nodes have to be computed. For this, we use the technique proposed in [26]. The numerical model is based on the finite elements method (FEM), using the algebraic subgrid scale (ASGS) as in [27] for stabilisation. In order to solve this system efficiently in supercomputers, a split approach is used [28]. The Schur complement is obtained and solved with an Orthomin(1) algorithm [29] with weak Uzawa operator preconditioner. The momentum equation is solved twice using generalized minimal residual method (GMRES) with Krylov dimension 100 and a diagonal preconditioner. For the continuity equation a deflated conjugate gradient algorithm is used. Further details of the heart model can be found in [30].

#### 2.2.3 Initial and boundary conditions

The Mitral inlet has a constant pressure of *P_LA_*. The aortic model has a 0D Windkessel model with three components: a resistor*R_s_*, serially connected with an *RC* parallel impedance *R_p_*, and *C_p_* [21]. On the deforming ventricular walls, as well as on the rest of the fluid domain, the velocity the domain deformation is imposed. This is *v_i_* |_Γ_*d*__ = *∂b_i_/∂t*, where *v_i_* |_Γ_*d*__ is the velocity at the deformed boundary. The LVAD outlet has a specific type of boundary condition that will be thoroughly described in Section 2.2.5. the rest of the boundaries have *v_i_* |_Γ_*r*__ = 0. As for the initial condition in the unknowns, the initial fluid velocity is *v_i_* |_*t*=0_= 0.

#### 2.2.4 Valve modelling

Mitral and aortic valves are modelled through a pressure-driven porous layer in the valvular region. This porous media add an isotropic force to the right-hand side of the momentum equation with the shape 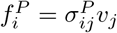, where 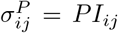 where *P* is the material porosity and *I_ij_* the identity matrix. This strategy provides a robust numerical scheme against the potential ill-conditioned stage of confined fluid. To ensure a smooth change with potentially abrupt changes in the transvalvular pressure, the porosity is driven through a hyperbolic tangent as:

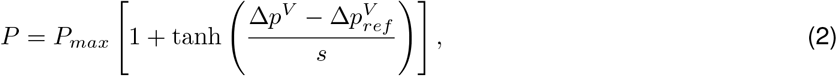

where *P_max_* the maximum possible porosity, *s* the slope of the curve, Δ*p^V^* the transvalvular pressure drop, and 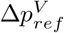 a reference pressure gradient. In practice if 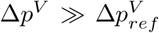 the valve is closed and if 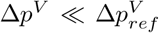 the valve is open. To avoid spurious valve opening and/or closing due to transient peaks in the tranvalvular pressure gradients, the measures are filtered using a median filter.

#### 2.2.5 LVAD boundary condition

Pump performance can be characterised with pressure-flow curves for each speed of the pump’s rotor. These pressure-flow curves (also called H-Q curves) provide a relation between the pressure difference between the pump inlet and outlet, and the flow the pump can provide at that speed.

Let Δ*p*^VAD^ = *P_Ao_* – *P_LV_* be the pressure difference between the outlet and the inlet of the pump and *Q*_VAD_ the flow through the pump. The pressure-flow relationship can be approximated with a quadratic equation as:

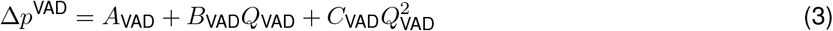

For each Δ*p*^VAD^ there is a single *Q*_VAD_ and vice versa. This relation can be used as a boundary condition, imposing a flowrate for a calculated pressure difference Δ*p*^VAD^. With this, the LVAD boundary condition keeps a flowrate constrained with Eq. (3).

#### 2.2.6 DARE

Executing the SA and the UQ during the VVUQ process requires generating thousands of inputs for the simulation code, submitting the jobs, processing the simulation results, and extracting the quantities of interest (QoIs) out of the physical field results. For this, we created DARE.

DARE is an automating tool that works coupled with Sandia’s Dakota [31, 32] and allows automatically encoding, submitting and retrieving jobs to any high performance computing (HPC) infrastructure (Fig. 3). Dakota allows to characterise and sample model inputs for multiple analysis types like SA, UQ or optimisation. The Dakota+DARE pair runs in a computer external to the HPC machine for as long as the analysis under execution may last (up to several weeks in this work). DARE receives Dakota’s chosen inputs, processes the simulation templates with an encoder, submits the job to the supercomputer queue, waits for the jobs to be finished and processes the final results to feed Dakota with the obtained outputs. A combination of Dakota’s restart capabilities with DARE failure capture capabilities makes this a robust framework for the required analysis.

**Fig. 3:**
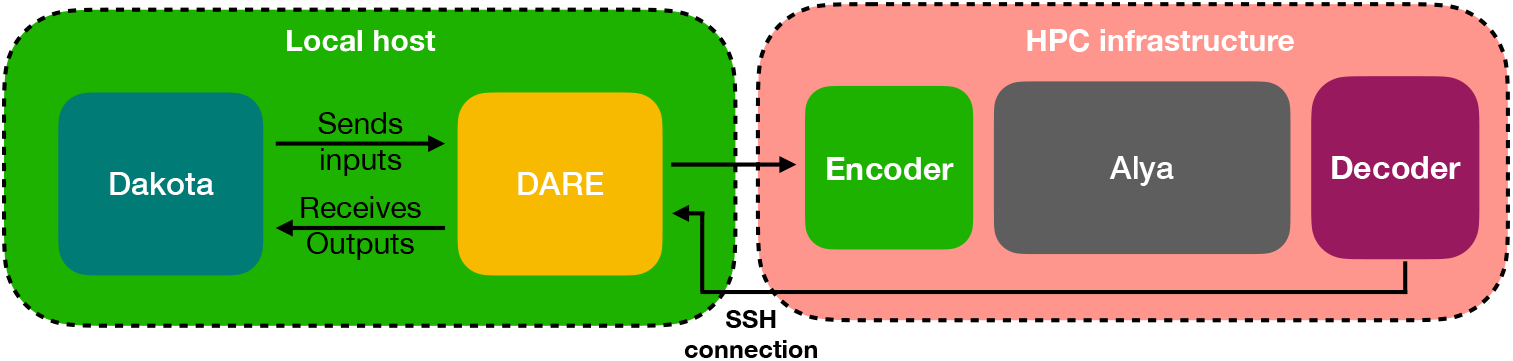
Scheme of DARE building blocks.

### 2.3 Design of the VVUQ plan through V&V40 risk-based credibility assessment

We performed credibility assessment by following the ASME V&V 40 [10] standard. The standard provides a framework for assessing the relevance and adequacy of the completed VVUQ activities for medical devices. Applying the standard requires a set of preliminary steps to determine the required level of credibility for the model. These preliminary steps are to identify: *(1)* the question of interest, this is the question the tool will find an answer to; *(2)* the context of use (CoU), this is the specific role and scope of the computational model; *(3)* the QoI, these are the simulation outputs relevant for the CoU; *(4)* the model influence, this is the contribution of the model in making a decision; *(5)* the decision consequence, this is the possibility that incorrect model results might lead to patient harm; and *(6)* model risk, which is based on model influence and decision consequence. Once these items are identified, goals for the credibility evidence can be defined and the VVUQ plan designed.

#### Question of Interest

For an apically implanted LVAD, does the selected pump speed produce: *(a)* complete aortic valve opening (*Q_Ao_* > 5[*cm*^3^/*s*]); and *(b)* a Cardiac output compatible with life 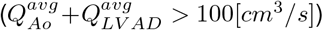 for a range of HR and EF covering a HF patient population?

#### Context of Use (CoU)

The heart-LVAD computational model may be used by design engineers to assist in the preclinical development of LVAD, by characterising aortic root, LVAD and intra-LV flows for a given pump speed. The goal of the heart-LVAD computational model is to provide a computational replica of a benchtop experiment for a quantitative analyses in parametric explorations. The heart-LVAD computational model by no means is replacing animal experiments or clinical trials, but augmenting the totality of evidence.

#### Quantities of Interest (QoI)

These are the simulation outputs relevant to the CoU. For completeness, here we repeat that the QoIs are the maximum and average flows through the outlet boundaries (LVAD flow *Q_V AD_* and aortic root flow *Q_Ao_*).

#### Model influence

Although the numerical test will augment the evidence provided by the bench test to aid design, they do not qualify the safeness of the device. This meaning animal testing and clinical trials are still required to prove safety and efficacy of the device. Therefore the computational model influence can be categorised as low (see Table 1).

**Table 1:** Risk map. Adapted from [33].

|  |  |  |  |  |
| --- | --- | --- | --- | --- |
| Model influence | high | 3 | 4 | 5 |
|  | med | 2 | 3 | 4 |
|  | low | 1 | 2 | 3 |
|  |  | low | med | high |
|  |  | Decision consequence |  |  |

#### Decision consequence

If the model fails to make accurate predictions for the question of interest, could advice for an operating condition that produce either: *(a)* low cardiac output or *b* a permanently closed aortic valve. These might lead to thromboembolic events, aortic regurgitation or death. Therefore the decision consequence is categorised as high (see Table 1).

#### Risk assessment

As the model influence has been categorised as “low” and the decision consequence as “high”, the LV-LVAD model is categorised with a risk of 3 on the 1-5 scale from Table 1, therefore requiring a medium level goals in the VVUQ plan.

##### 2.3.1 Translation of model risks into credibility goals

The ASME V&V40 standard [10] defines 13 credibility factors (some containing sub-factors) that break down the assessment of the VVUQ activities. Once the risk associated with the modelling tool has been determined, the next stage of the V&V40 pipeline is defining a ranking (gradation) for each factor sorted by increasing level of investigation, and then selecting a credibility goal for each factor based.

Table 2 lists the 13 credibility factors and sub-factors. Gradations for each factor are provided in the supporting material, Section S1. For most credibility factors, the gradation proposed in the V&V40 standard is used. Table 2 summarizes the maximum possible score in the gradation, the targeted goal and the achieved score. The targeted goal also includes the description required to achieve that score. Per V&V40, goals were chosen so that model credibility is generally commensurate with model risk. Therefore, for most factors, a medium level or higher goal was chosen. The rationale behind the chosen goals is provided in the supporting material, Section S2.

**Table 2:**
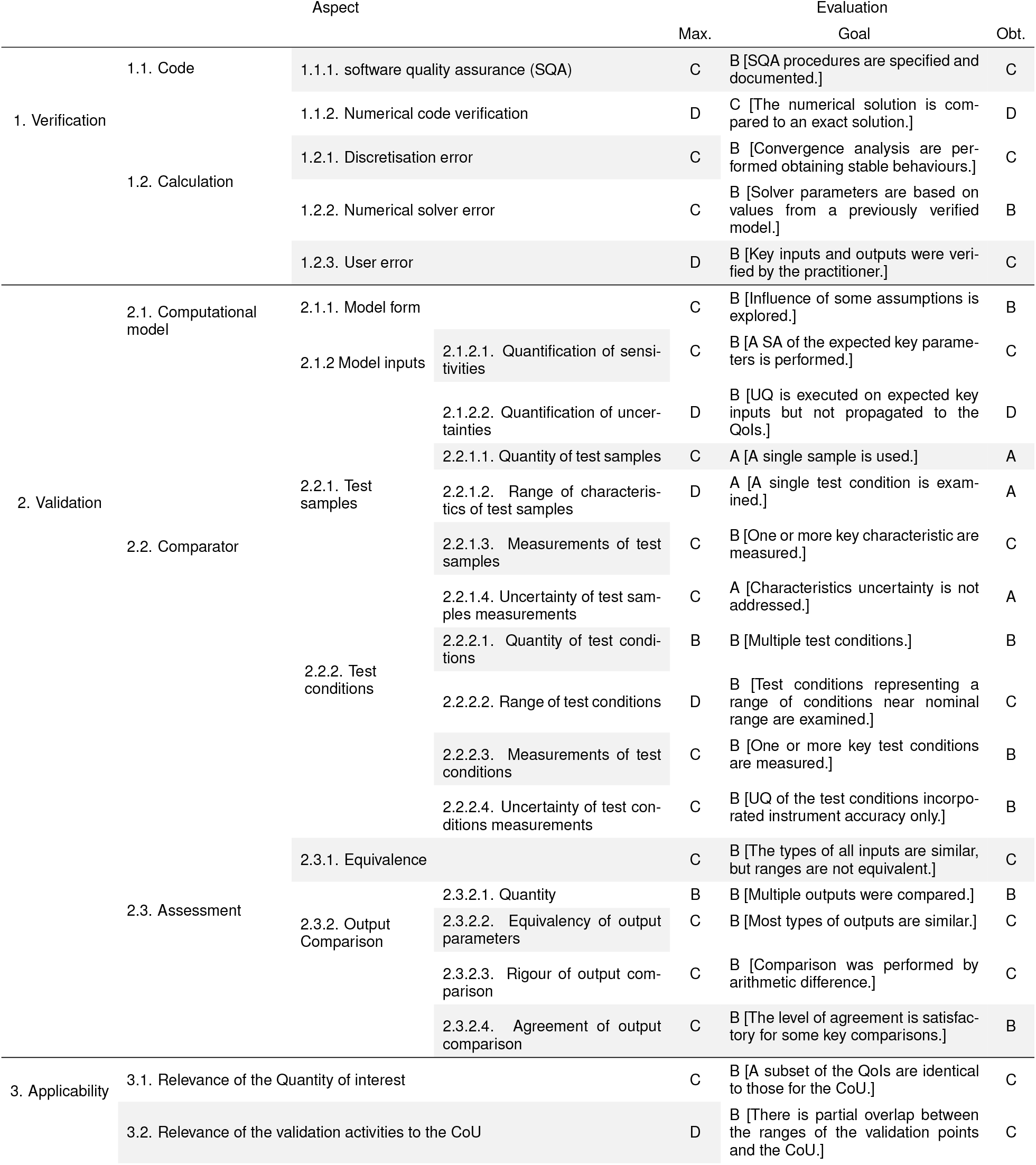
ASME V&V40 credibility factors [10] analysed on the risk-based assessment. The table shows the maximum possible score (“Max.” column), the desired goal (“Goal” column) and the obtained score (“Obt.” column). The goal column also includes the description of the activity to achieve that gradation.

### 2.4 Design and goals of the VVUQ plan

This section explains the VVUQ activities carried out to achieve the credibility goals defined in Section 2.3.1. The VVUQ plan has been designed following [10, 11, 8].

#### 2.4.1 Steps of the VVUQ plan

1. **Provide verification evidence:** SQA practices should be followed to ensure reproducibility and traceability. Numerical code verification (NCV) is mandatory to ensure correctness in the coding of the models. Numerical calculation verification is mandatory to ensure a sufficient spatial discretisation of the problem.
2. **Execute a sensitivity analysis in the operating range:** A non-linear global SA within the operating range of the cases should be executed to: *(a)* understand the impact of each input on the QoIs, and *(b)* Safely reduce the number of input variables for the UQ through Pearson’s ρ and Sobol indices analyses. The goal of step *(b)* has a direct impact in the UQ as it reduces its computational cost.
3. **Perform validation with uncertainty quantification:** The reduced input model obtained from the SA is used to execute the UQ analysis. At least a middle point and the extreme cases of the operation envelope should be investigated. A comparison of the QoIs’ distributions is required including a validation metric that allows quantitatively comparing the results between validation points and against other similar works or future projects.
4. **Adequacy assessment:** Evaluate if the simulation credibility evidence is good enough to safely answer the question of interest.

For the sake of brevity, the calculation and code verification tests are omitted in this text and shown in the supporting material, Section 3. The rest of the present manuscript focuses on the SA, validation, and UQ.

#### 2.4.2 Sensitivity analysis

A SA [34] is a statistical tool that allows quantifying the impact of each input variable in each QoI of the model. It is helpful as it allows to rank the input variables based on their contribution to the variation of the model output. On the one hand, identifying the less relevant inputs allows to reduce the dimensionality of the problem, as the less relevant inputs can be safely avoided to decrese the computational cost during UQ. On the other hand, reducing the experimental uncertainty of the most relevant inputs identified is critical to obtain accurate model predictions. The reason for this is that a large uncertainty in a highly impactful input will produce a large uncertainty in the model output.

In this work we execute two types of SA: *i)* local SA via Pearson coefficients [35], and *ii)* global SA through total Sobol indices calculation [36] by relying on a 5-th order polynomial chaos expansion (PCE) [37]. Both the analysis are performed by relying on 500 samples. However, while the former is a measure of the strength of a linear association between an input and is valid under the assumption of linear and homoskedastic data with no multivariate outlayers, the latter provides information of the importance of each input taking into account complex factors like nonlinearities, input interactions, and sample dispersion.

#### 2.4.3 Uncertainty quantification

Validation involves measuring the difference between both sources of predictions, the experiment and the simulation. These two sources are subject to different types of uncertainties that should be identified as part of a UQ analysis.

The measuring instruments in the experiment introduce the measuring error, while user error is introduced by the experimentalist variability. In the experiment on this manuscript, each execution of the experiment contains multiple beats. Therefore, even for the same set of inputs and due to the measuring instrument error (see Section 2.1), there will be a dispersion in the QoIs that requires the the measurement error to be quantified. As we use retrospective experimental data not specifically thought for VVUQ, there is only a single execution of the experiment for each of the six validation points. This hampers quantification of the user error. To tackle this issue we add a 10% error range in the QoIs measured in the experiment to account for the user error. As there is no other information on that user error shape, no probability distribution can be assumed. The numerical error in the simulation tool is estimated by the code and calculation verification (see supporting material Section S3), and the input error quantified during the UQ. The simulation input variables are listed and classified in Table 3.

**Table 3:** Table of model inputs and their uncertainty characterisation.

| Variable | Symbol | Uncertainty classification | Characterisation | value range |
| --- | --- | --- | --- | --- |
| Density | $\rho$ | deterministic | exact value | $1.1[g/cm^2]$ |
| Viscosity | $\mu$ | deterministic | exact value | $0.0372[Poise]$ |
| Heart rate | $HR$ | aleatory | uniform | $[40, 120][bpm]$ |
| Atrial pressure | $P_{LA}$ | aleatory | uniform | $[5, 15] \times 1 \times 10^3[Baryes]$ |
| Ejection fraction | $EF$ | aleatory | uniform | $[0.1, 0.35][\%]$ |
| Aortic serial resistance | $R_s^{Ao}$ | aleatory | uniform | $[50, 200][\frac{Baryes \cdot s}{cm^3}]$ |
| Aortic parallel resistnace | $R_p^{Ao}$ | aleatory | uniform | $[0.5, 2] \times 1 \times 10^3[\frac{Baryes \cdot s}{cm^3}]$ |
| Aortic parallel Capacitance | $C_p^{Ao}$ | aleatory | uniform | $-[0.5, 2] \times 1 \times 10^{-3}[\frac{cm^3}{Baryes}]$ |
| Constant VAD coefficient | $A_{VAD}$ | aleatory | uniform | $[25.3, 1] \times 1 \times 10^6[\frac{cm^3}{s}]$ |
| Linear VAD coefficient | $B_{VAD}$ | aleatory | uniform | $[0.5, 5] \times 1 \times 10^3[\frac{cm^3 \cdot Baryes}{s}]$ |

Each one of the model variables can be characterised as one of the following three: *(a)* deterministic, when their values are known *(b)* aleatory, when the variable is uncertain due to inherent variation and can be characterised with a cumulative distribution function (CDF); *(c)* epistemic, when the variable is affected by reducible uncertainty due to lack of knowledge and it can be represented with a bounded interval. The inputs of the model should be characterised by one of these three categories and treated accordingly. These identified uncertainties are forward propagated through the computational model down to the output to obtain the QoIs distributuons. Once the output distributions are obtained for the experiments and for the simulations, the differences are quantified using a validation metric.To evaluate these differences, we use the Minkowski *L*_1_ norm (MN) validation metric proposed in [8] in two different ways. For the first approach (called MN^*u*^) a uniform distribution is assumed in the experimental data, and the MN integrated between that artificially built experimental empirical cumulative distribution function (ECDF) and the simulation ECDF. For the second approach, so called p-box approach [38], no distribution is assumed in the experimental data. Therefore, the MN is calculated between the simulation ECDF and the maximum (MN^+^) and minimum (MN^−^) limits of the experimental data.

## 3 Results

Results are split in two. Section 3.1 shows the results for the SA of the numerical model, which rank the model input variables according to their impact on the outputs. The SA is carried out by firstly sampling the input values using latin hypercube sampling (LHS) for all the input variables. Later, these samples are used to execute independent CFD simulations. Results are analysed through scatter plots, Person’s ρ correlation and total Sobol indices. Section 3.2 shows a UQ analysis for six validation points reproduced in the SDSU-CS. The six validation points include two different conditions, so called 22[%]@68.42[*bpm*] and 17[%]@61.18[*bpm*], with three pump speeds (0*k*, 8*k* and 11*k*[*rpm*]) each.

The model is intended to reproduce inbound and outbound flows in the LV of the CS, therefore the final goal is to correctly reproduce the flow meter signals of the experiment. As the statistical tools for the UQ analysis require scalars, the QoIs chosen to characterise the flows are maximum and average Aortic and LVAD flows.

An analysis and comparison of each set of simulation results at every spatial point of the volumetric domain is virtually impossible even for a small number of cases, let alone more than 1000 simulation executions as done in this work. Therefore, even if sample qualitative results are shown for each validation point, the maximum and average flows through the boundaries are calculated and used to calculate statistical trends.

### 3.1 Sensitivity Analysis

#### 3.1.1 Results for the SA

The SA is intended to identify the input variables with the highest impact in the QoIs. The variable ranges used for the SA are shown in Table 3.

The density *ρ* and the dynamic viscosity *μ* are easily and accurately measured. Furthermore, due to the system operation pressures and the fluid bulk properties, these quantities are not expected to change. With this, these two variables are classified as deterministic, knowing their exact value. The rest of the input variables are ranged in approximately one order of magnitude. To proceed with the LHS a uniform distribution is considered, obtaining 500 samples from the input variables. These samples are used to run 500 simulations, obtaining an ensemble of the QoIs. The sampling and results are shown in the scatter plot at Fig. 4 together with the Person’s correlation coefficient ρ. From a visual analysis of the scatter plot it can be seen that the data is nonlinear, heteroskedastically distributed, and contains multivariate outliers, failing 3 of the 7 assumptions required for Pearson’s analysis. To overcome this issue, a global SA is done by calculating total Sobol indices (indicated in Section 2.4.2). Total Sobol indices provide information of the importance of each input taking into account complex factors like nonlinearities, input interactions, and sample dispersion. The total Sobol index of each input with respect to each QoI are shown as a tornado plot in Fig. 4. The larger the index, the more important that input is for the QoI. The total cost of the 500 simulations is about 50 [core-years] in Marenostrum IV supercomputer.

**Fig. 4:**
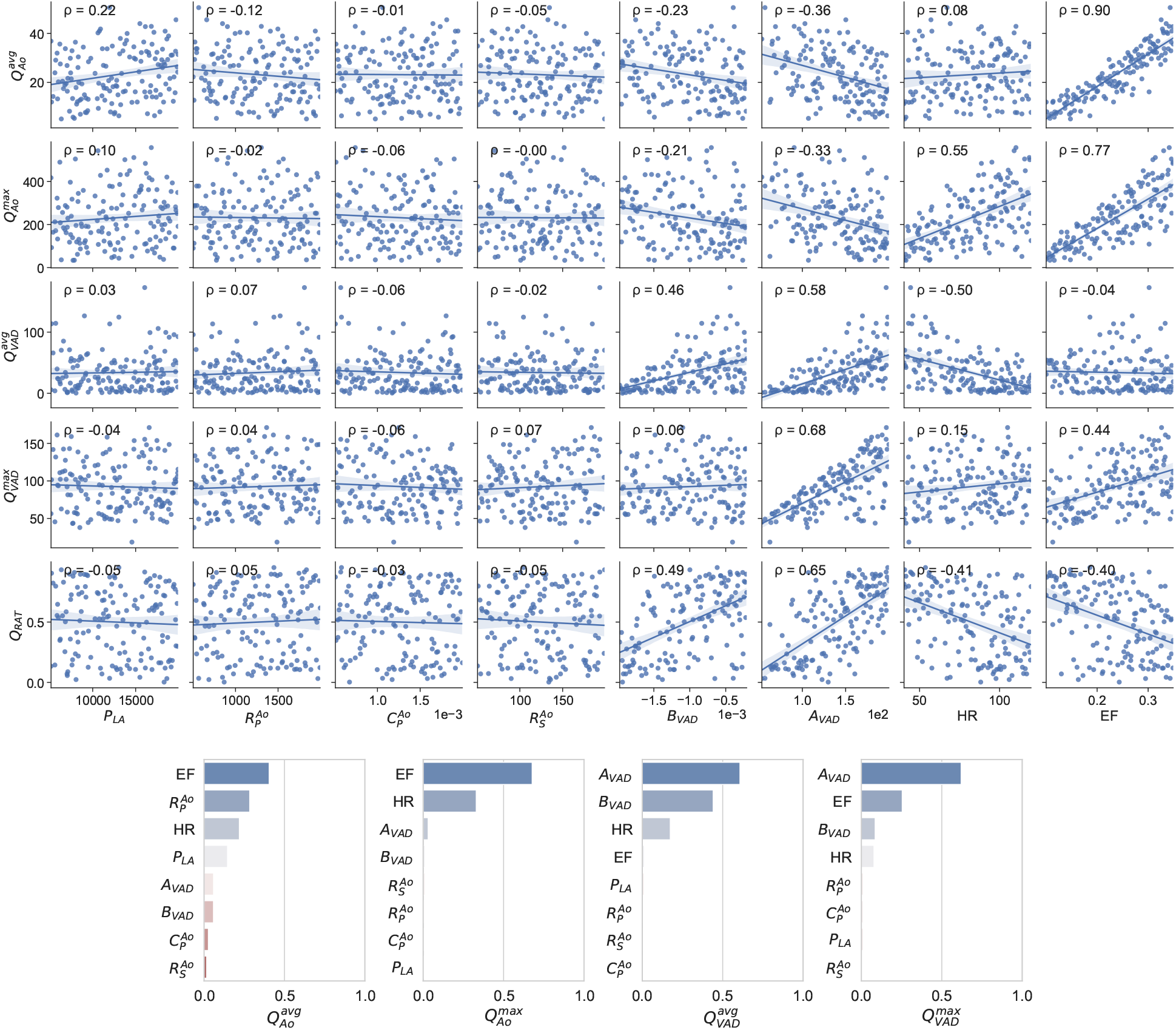
Scatter plots and total Sobol indices tornado plots for the 8 input variables and the 6 QoIs. The scatter plot also shows the Pearson’s linear correlation number ρ in the top left corner.

#### 3.1.2 Discussion of the SA results

The scatter plots and the Pearson’s ρ analysis shown in Fig. 4 provide a simple tool to identify the most important variables in the LV-LVAD system. While these tools are useful for a first order approach to understanding the system’s behaviour, they fall apart for non-linear effects and complex interactions. Total Sobol indices provide a more insightful tool that account for the effect of each input variable in each QoI. From the total Sobol indices analysis we can see that the most highly ranked inputs are the EF, HR, *A_V AD_* and *B_V AD_*. Finally, the wide ranges chosen for the global SA provide a trustworthy set of Sobol indexes that are applicable to the smaller ranges during the UQ analysis.

While SA is a common tool other fields of cardiac modelling like electrophysiology or solid mechanics [39, 40, 41], there is no published work with a local nor a global SA for ventricular CFD. Despite this, The trends in Fig. 4 agree with experiment data and clinical observations. A higher pump speed, translated as a larger *A_V AD_* coefficient, has a positive correlation with the LVAD flow and a negative correlation with the aortic flow [42]. The reason for this is that the suction produced by the pump reduces the aortic valve opening [43, 7]. Also, the HR and EF has a direct positive correlation with the LVAD and aortic flows [44]. The unexpectedly [45, 44] small influence of the mean atrial pressure (*P_LA_*) and arterial impedance (characterised via 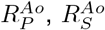 and 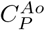) can be explained due to the lack of Frank-Starling mechanism [46] in the silicone ventricle of the experiment and therefore also in its computational analogue.

### 3.2 Validation with uncertainty quantification

The global SA in Section 3.1 identified the four most relevant variables, namely the EF, the HR, *A_V AD_*, and *B_V AD_*. These are the simulation input variables studied during the UQ analysis. The UQ analysis consist of six validation experiments varying the four chosen inputs. For each validation point, a qualitative set of images is shown that allows visualising the CFD behaviour of the problem. The quantitative results are analysed through scatter plots and ECDFs. To evaluate the differences between the experimental and simulation distributions, we use the MN validation metric already described in Section 2.4.3. In every figure, experimental results are represented with orange and simulation results with blue.

#### 3.2.1 Validation points and ranges

For the UQ analysis, the SDSU-CS (Section 2.1) is configured at two beating conditions. The condition 22[%]@68.42[*bpm*] has an EF= 22[%] and HR= 68.42[*bpm*]. The condition 17[%]@61.18[*bpm*] has an EF= 17[%] and HR= 61.18[*bpm*]. Three pump combinations are used in each case, 0*k*[*rpm*] (or pump off with clamped outflow LVAD conduit, recall Section 2.1), 8*k*[*rpm*], and 11*k*[*rpm*]. This makes a total of six validation points as described in Table 4. As there is no information on the precision of the prescribed EF and HR, a 10% error is assumed for these two inputs, producing the ranges in the second and third column of Table 4. Similarly, and as explained in Section 2.4, the experiment data accounts for the instrument error. But, as we count with only a single experiment per validation point, an error range of 10% is included in the QoIs to account for the measurement uncertainty. To calculate one of the validation metrics shown (MN^*u*^) we assume a uniform distribution in the measured QoIs for that assumed range. On the contrary, the multiple simulations executed let us calculate the ECDF used for the metrics.

**Table 4:** The six validation points used for the UQ analysis.

| Condition | EF value range | HR value range | Pump speed |
| --- | --- | --- | --- |
| 22[%]@68.42[bpm] | [19.8,24.2][%] | [65.55, 72.45][bpm] | 0k[rpm] |
|  |  |  | 8k[rpm] |
|  |  |  | 11k[rpm] |
| 17[%]@61.18[bpm] | [15.3,18.7][%] | [57,63][bpm] | 0k[rpm] |
|  |  |  | 8k[rpm] |
|  |  |  | 11k[rpm] |

**Table 5:** Fitting coefficients (*a_V AD_, b_V AD_*, and *c_V AD_*) and fitting errors *ϵ* of the H-Q performance curves. The pump speed is measured in [*rpm*]. The units are *a_V AD_*[*Baryes*], *b_V AD_*[*Baryes · s/cm*^3^], and *c_V AD_*[*Baryes · s*^2^/*cm*^6^].

| Pump speed | $a_{VAD} \pm \epsilon / b_{VAD} \pm \epsilon / c_{VAD}$ |
| --- | --- |
| 0k | $0.0 \pm 0.0 / 0.0 \pm 0.0 / 0.0$ |
| 8k | $11.78 \times 1 \times 10^4 \pm 14.40 \times 1 \times 10^2 / -772.05 \pm 28.58 / 0.0$ |
| 11k | $21.71 \times 1 \times 10^4 \pm 48.90 \times 1 \times 10^2 / -902.81 \pm 61.55 / 0.0$ |

The method to calculate the variable ranges for the pump model inputs is explained in Appendix A. The coefficients range and the uncertainty characterisation are shown in Table 6.

**Table 6:** Range of H-Q curve coefficients used for the UQ analysis. The range is obtained as 2*ϵ_num_* + *ϵ_fit_* where *ϵ_fit_* is 10% of the measured value.

| Pump speed | coefficient | Uncertainty classification | characterisation | value range |
| --- | --- | --- | --- | --- |
| 0k | $a_{VAD}$ | deterministic | exact value | 0.0 [Baryes] |
| | $c_{VAD}$ | deterministic | exact value | 0.0 [Baryes · s<sup>2</sup>/cm<sup>6</sup>] |
| | $c_{VAD}$ | deterministic | exact value | 0.0 [Baryes · s<sup>2</sup>/cm<sup>6</sup>] |
| 8k | $a_{VAD}$ | aleatory | uniform | $[10.91, 12.66] \times 1 \times 10^4$ [Baryes] |
| | $b_{VAD}$ | aleatory | uniform | $-[7.90, 7.53] \times 1 \times 10^2$ [Baryes · s/cm<sup>3</sup>] |
| | $c_{VAD}$ | deterministic | exact value | 0.0 [Baryes · s<sup>2</sup>/cm<sup>6</sup>] |
| 11k | $a_{VAD}$ | aleatory | uniform | $[19.64, 23.77] \times 1 \times 10^4$ [Baryes] |
| | $b_{VAD}$ | aleatory | uniform | $-[9.80, 8.24] \times 1 \times 10^2$ [Baryes · s/cm<sup>3</sup>] |
| | $c_{VAD}$ | deterministic | exact value | 0.0 [Baryes · s<sup>2</sup>/cm<sup>6</sup>] |

These simulation variables are sampled using a LHS obtaining 50 samples per validation experiment, making a total of 300 numerical simulations that required about 30 [core-years] in Marenostrum IV supercomputer.

#### 3.2.2 Condition 22[%]@68.42[*bpm*]

Figure 5 shows qualitative surface results for the three pump speeds for the condition 22[%]@68.42[*bpm*] and the pump speeds 0*k*, 8*k*, 11*k*[*rpm*]. Afterwards, Figs. 6 to 8 provide quantitative results for the referenced condition and pump speeds. The flow plots Figs. 6a, 7a and 8a show the aortic (*Q_Ao_*) and LVAD flow (*Q_V AD_*) for the experiment (orange) and the simulation (blue). As the UQ analysis also accounts for HR, the time axis is normalised. The scatter plots Figs. 6c, 7c and 8c show the inputs in the x-axis and the outputs in the y-axis for the experiment (orange) and the simulation (blue). The orange range in the y-axis is representing the assumed 10% measurement error and the simulation results kernel distribution estimation (KDE) is represented in blue shades surrounding the simulation measures. Figures 6d, 7d and 8d show the simulation ECDF in blue and the experimental limits with two vertical orange ranges, together with the artificial uniform distribution used to calculate MN^*u*^. Finally, the bench and numerical experiments data limits and the multiple MN are shown in Figs. 6b, 7b and 8b.

**Fig. 5:**
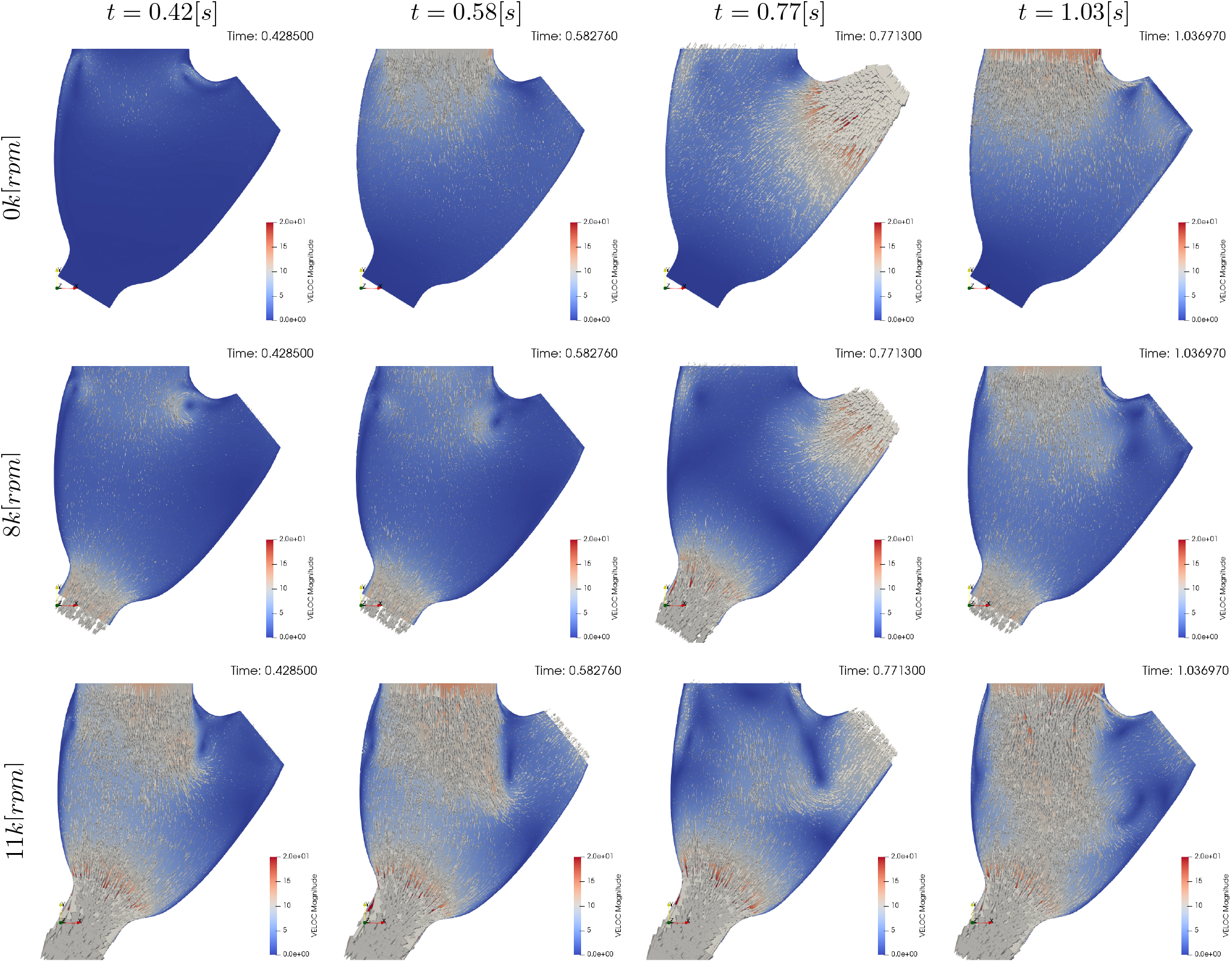
Qualitative surface results for the condition 22[%]@68.42[*bpm*]. Pump speed is 0*k*[*rpm*], 8*k*[*rpm*], and 11*k*[*rpm*] in the first, second, and third rows respectively. The columns indicate different time frames in the simulation. *t* = 0.42[*s*] show results for the plateau previous to systole, *t* = 0.58[*s*] show the atrial kick, *t* = 0.77[*s*] show systole, and *t* = 1.03[*s*] shows diastole. Videos of the simulations can be found in the Supporting Material Video 1.

**Fig. 6:**
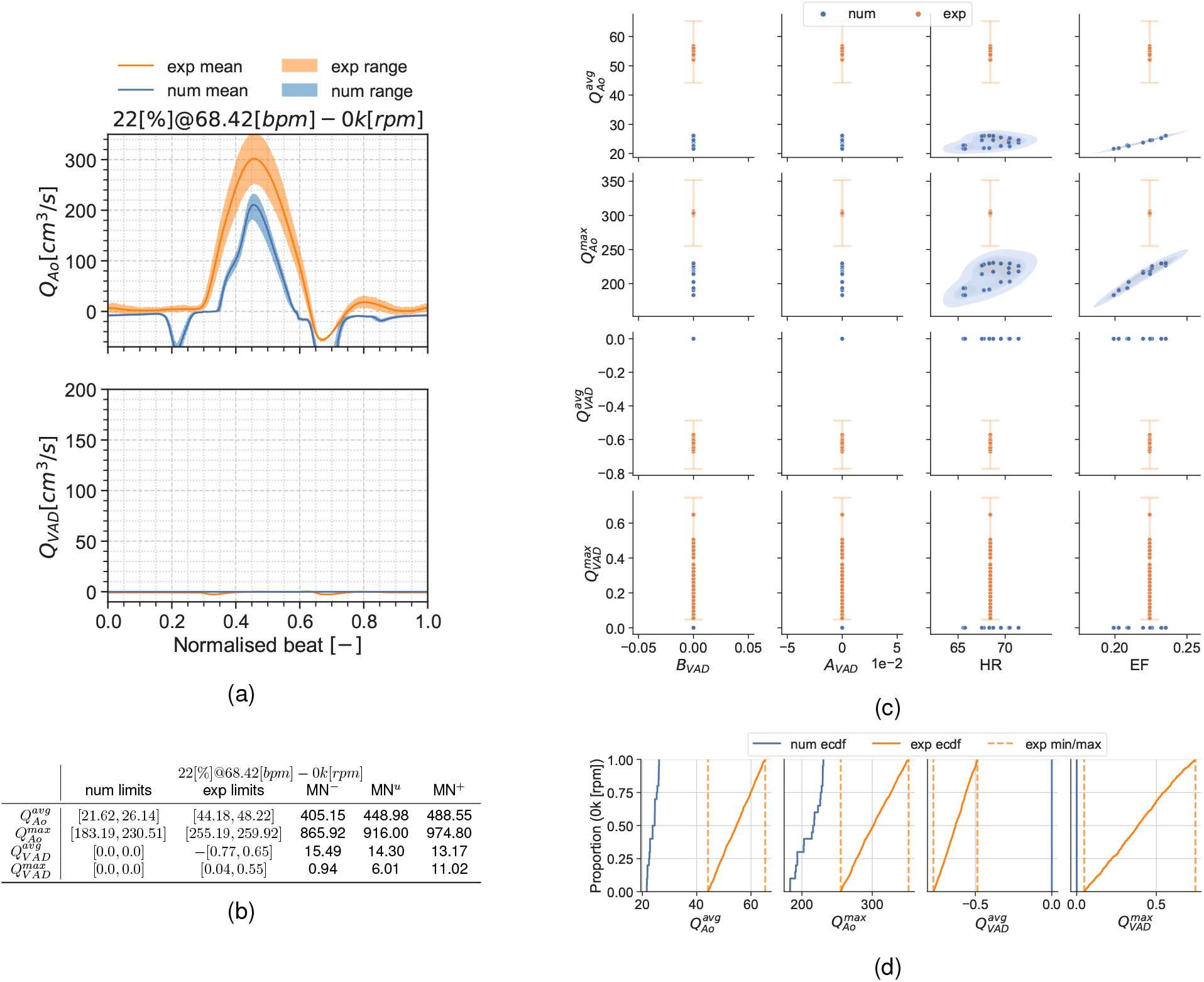
Summary for the condition 22[%]@68.42[*bpm*] and 0*k*[*rpm*]. 6a: aortic valve and LVAD flows. 6b: validation metrics. 6c: scatter plot showing the experimental and simulation data. 6d: ECDF for the simulation, experimental data limits and the constructed uniform distribution.

**Fig. 7:**
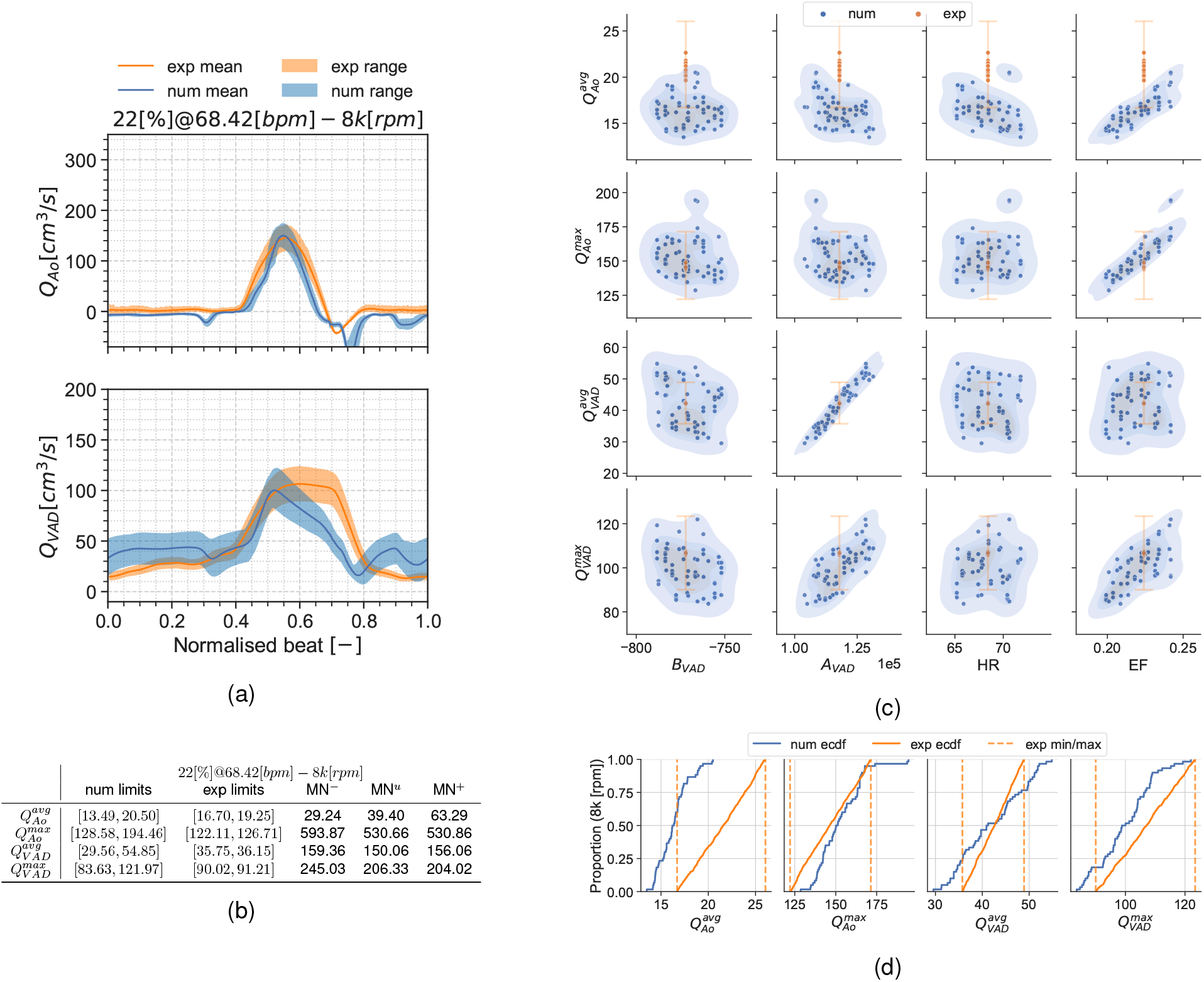
Summary for the condition 22[%]@68.42[*bpm*] and 8*k*[*rpm*]. 7a: aortic valve and LVAD flows. 7b: validation metrics. 7c: scatter plot showing the simulation and experimental data. 7d: ECDF for the simulation, experimental data limits and the constructed uniform distribution.

**Fig. 8:**
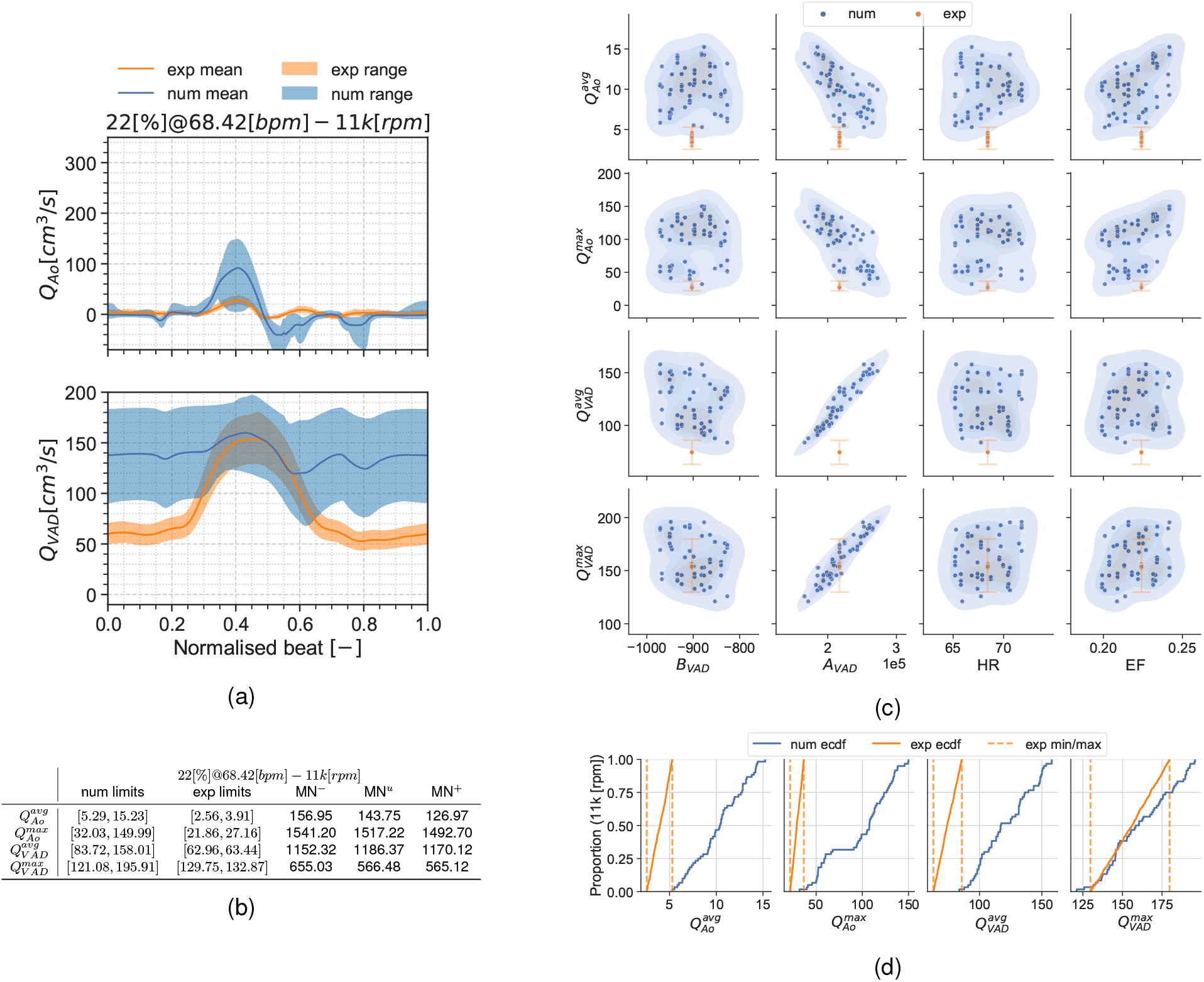
Summary for the condition 22[%]@68.42[*bpm*] and 11*k*[*rpm*]. 8a: aortic valve and LVAD flows. 8b: validation metrics. 8c: scatter plot showing the simulation and experimental data. 8d: ECDF for the simulation, experimental data limits and the constructed uniform distribution.

#### 3.2.3 Condition 17[%]@61.18[*bpm*]

Results are presented similarly to Section 3.2.2. Figure 9 show a set of time frames for the three pump speeds during the condition 17[%]@61.18[*bpm*]. Afterwards, Figures 10 to 12 show the experimental and simulation results. Again, the experimental results are shown in orange and simulation results in blue. the experiment and simulation ranges and the validation metric are shown in Figs. 10b, 11b and 12b.

**Fig. 9:**
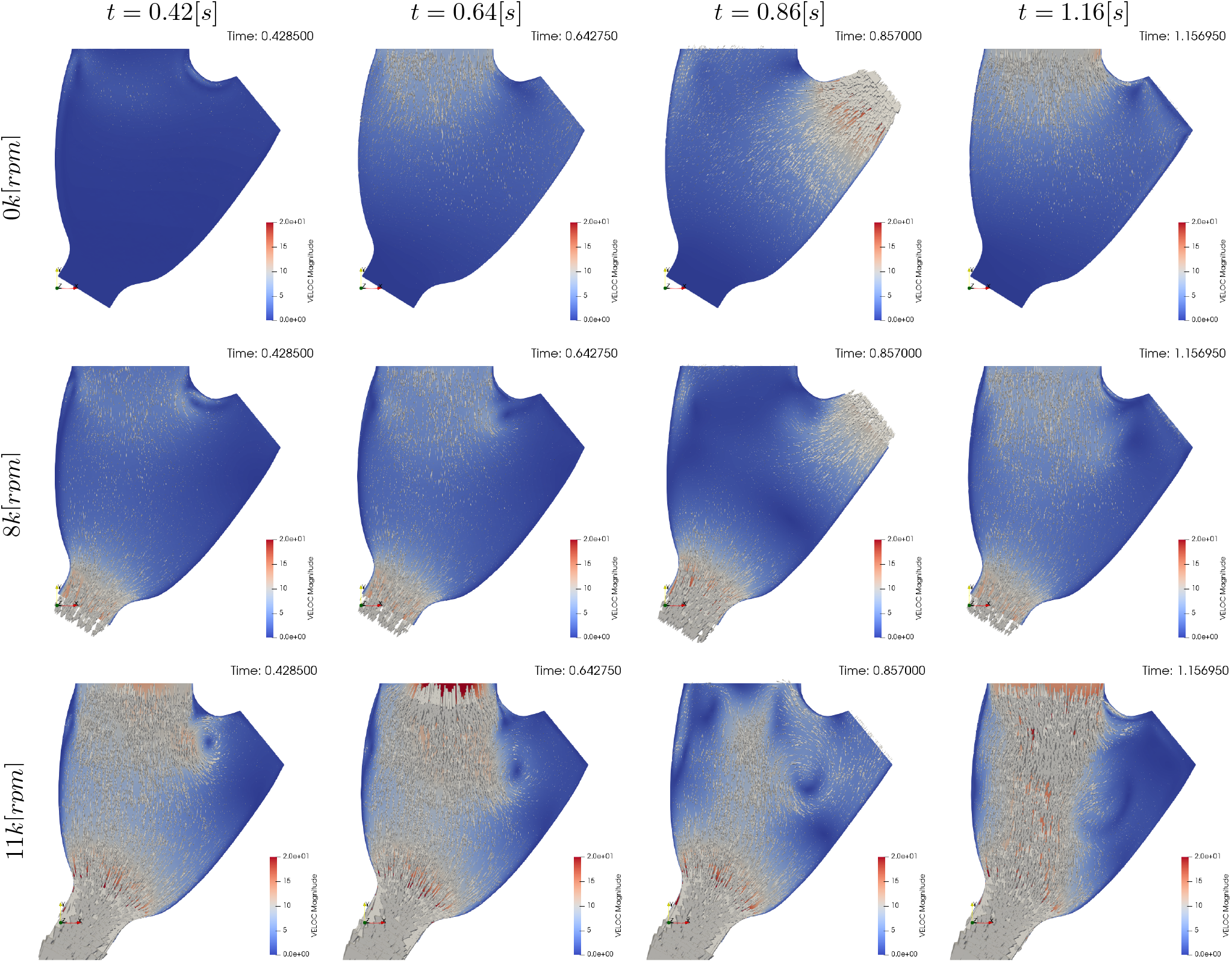
Qualitative surface results for the condition 17[%]@61.18[*bpm*]. Pump speed is 0*k*[*rpm*], 8*k*[*rpm*], and 11*k*[*rpm*] in the first, second, and third rows respectively. The columns indicate different time frames in the simulation. *t* = 0.42[*s*] show results for the plateau previous to systole, *t* = 0.64[*s*] show the atrial kick, *t* = 0.85[*s*] show systole, and *t* = 1.16[*s*] shows the diastolic filling. Videos of the simulations can be found in the Supporting Material Video 1.

**Fig. 10:**
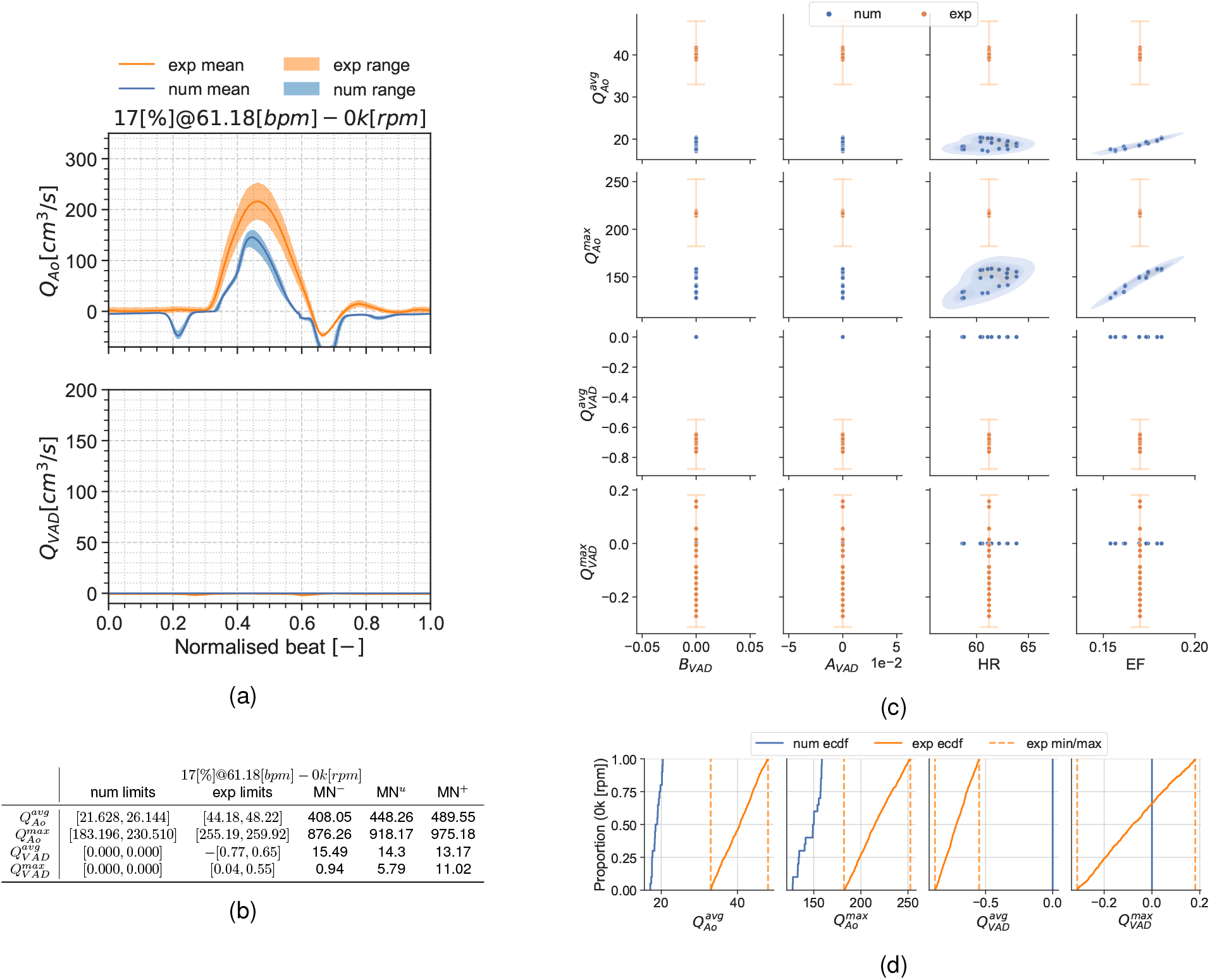
Summary for the condition 17[%]@61.18[*bpm*] and 0*k*[*rpm*]. 10a: aortic valve and LVAD flows. 10b: validation metrics. 10c: scatter plot showing the simulation and experimental data. 10d: ECDF for the simulation, experimental data limits and the constructed uniform distributions.

**Fig. 11:**
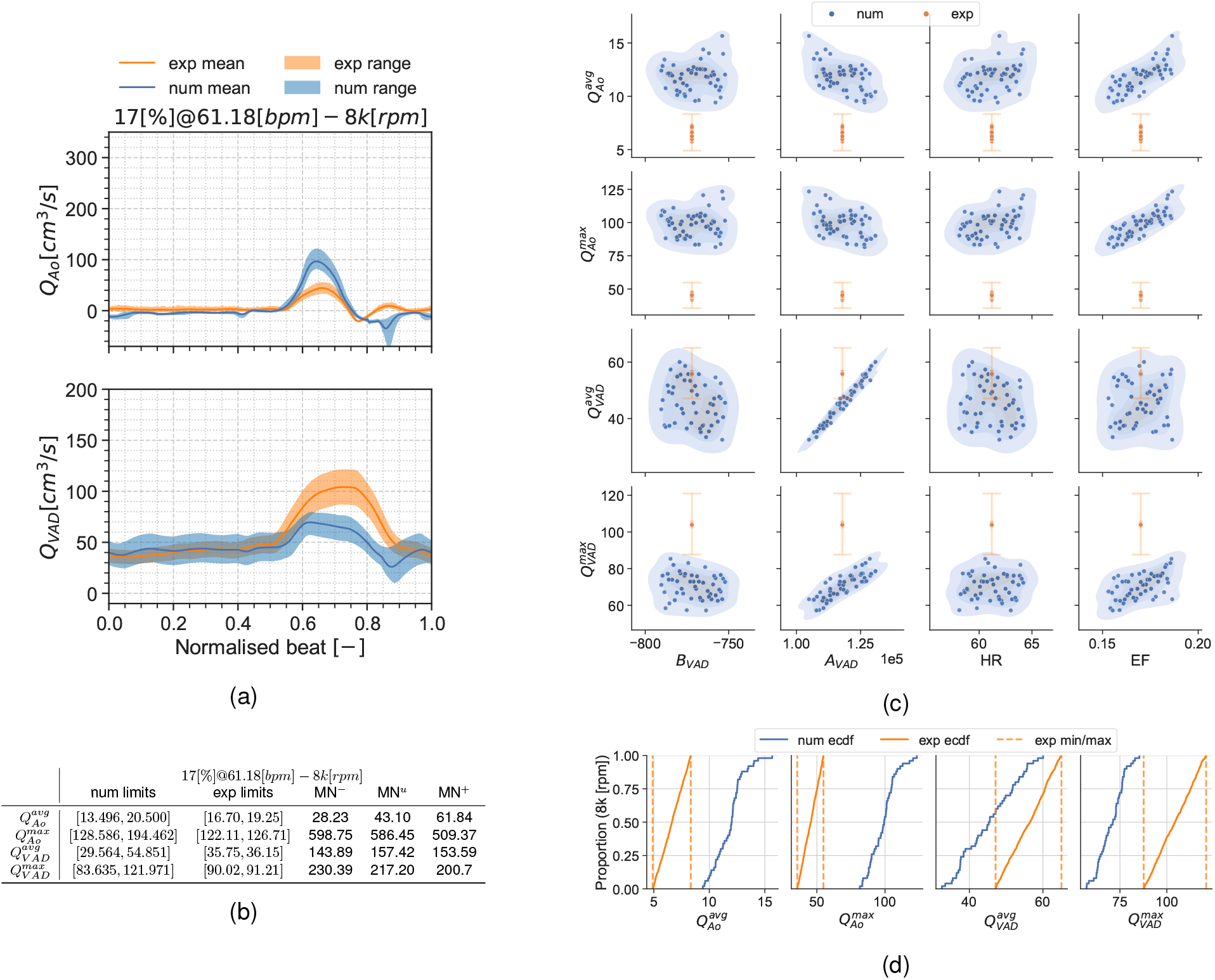
Summary for the condition 17[%]@61.18[*bpm*] and 8*k*[*rpm*]. 11a: aortic valve and LVAD flows. 11b: validation metrics. 11c: scatter plot showing the simulation and experimental data. 11d: ECDF for the simulation, experimental data limits and the constructed uniform distribution.

**Fig. 12:**
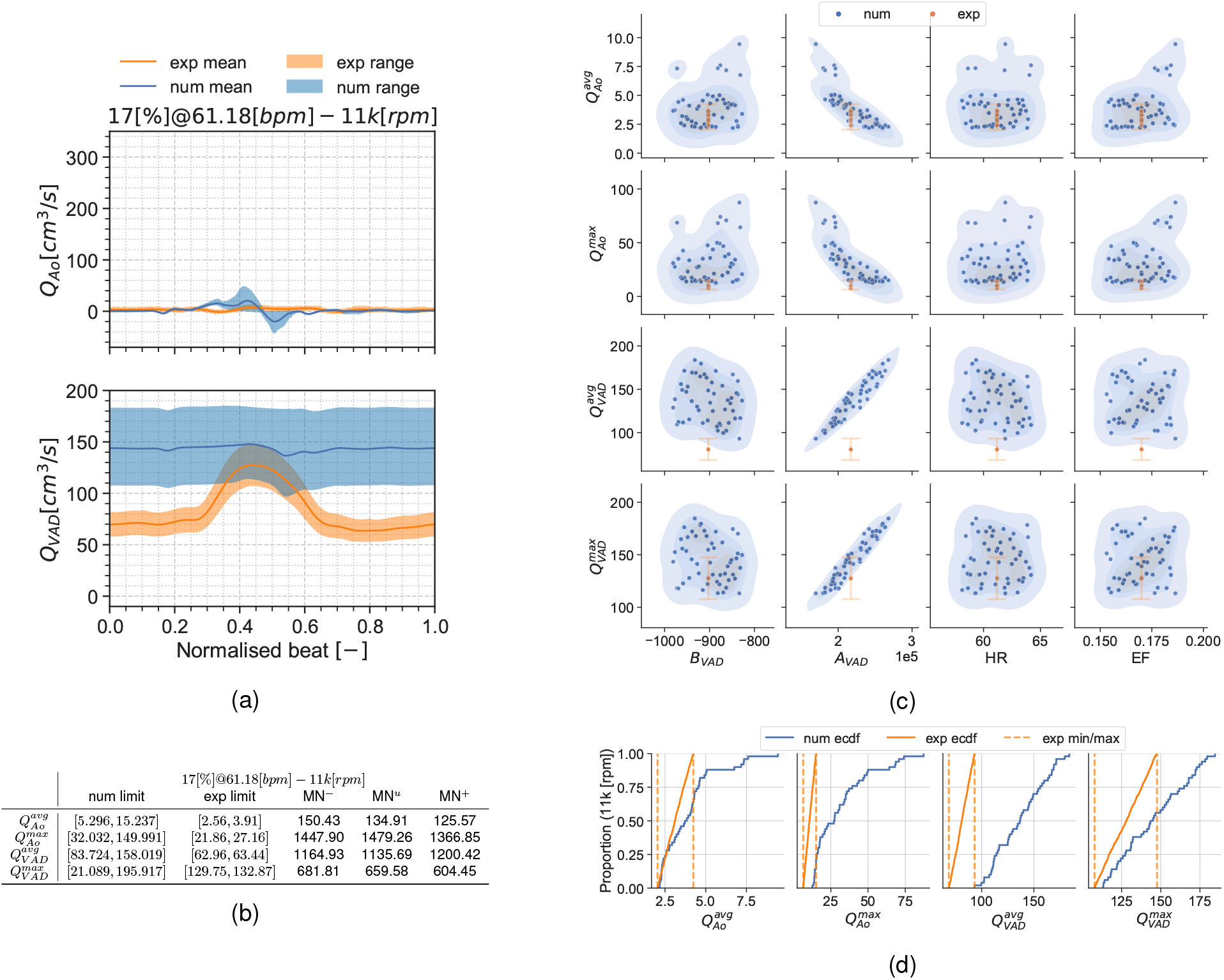
Summary for the condition 17[%]@61.18[*bpm*] and 11*k*[*rpm*]. 12a: aortic valve and LVAD flows. 12b: validation metrics. 12c: scatter plot showing the simulation aned experimental data. 12d: ECDF for the simulation, experimental data limits and the constructed uniform distributions.

#### 3.2.4 Discussion of the UQ results

Similarly to the SA, while UQ analyses have been recently done for electrophysiology and electromechanical models of the heart [41, 39, 47], but no equivalent studies have been conducted for ventricular CFD.

The first noticeable feature in the scatter plots is the lack of dispersion in the x-axes for the experimental results. The x-axes represent the prescribed values of the inputs in the experiment or simulation. As there is a single execution of the experiment per validation point, there is no information on the input variable distribution. This fact that translates as zero dispersion in the x-axis at experimental scatter plots. On the contrary, the multiple beats contained in each validation point and the measurement error in the flow meter is seen as y-axis dispersion for the experiment.

Another noteworthy detail is the fact that the MN is an absolute metric, therefore its interpretation depends on the QoI’s range and mean value at the specific condition. As an example, the 0*k*[*rpm*] cases for both conditions may seem the trivial solution for 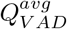 and 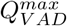. but, even if they have the smallest validation metrics in this manuscript, there is no overlap for these QoIs in the scatter plots.

Results show the smallest validation metrics (i.e. better agreement) for the mid-point working conditions with larger differences for the extreme cases. The large uncertainty ranges in the pump H-Q curve Fig. 14 and Table 6 for the 8*k*[*rpm*] and most noticeably 11*k*[*rpm*] speeds produce a considerable dispersion in the simulation results. This dispersion is most noticeable in the 11*k*[*rpm*] cases (Figs. 8 and 12) as a large uncertainty range in the flow curves (Figs. 8a and 12a), but it also exhibits itself in the ECDFs (Figs. 8d and 12d) and the scatter plots (Figs. 8c and 12c) as a poor overlap of the blue-shaded simulation KDE and the orange range for the experiments. Particularly, Figure 12c shows a clustering in the 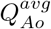 and 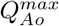 simulation points. If the aortic valve flow *Q_Ao_* is plotted for each execution (not shown in this manuscript) the clustering stands out as a set of curves where the Aortic valve fully opens, allowing flow through it. This is because the *a_V AD_* and *b_V AD_* uncertainty ranges are so wide that they even allow aortic valve opening for some scenarios, something also observed in [42]. The largest validation metric (Figs. 8b and 12b) with the pump operating at 11*k*[*rpm*] is for 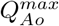 for both conditions 17[%]@61.18[*bpm*] and 22[%]@68.42[*bpm*].

The 0*k*[*rpm*] results at Figs. 6 and 10 also show a mismatch between the numerical and the bench data. As the pump H-Q curves are forced to zero in the simulation, there is also zero uncertainty in the associated inputs (*a_V AD_* and *b_V AD_*). This produces unequivocally LVAD flows equal to zero 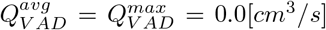 and a Dirac’s delta probability distribution function for these QoIs in the simulation results. On the contrary, the experiment still shows a y-axis scattering in the QoIs (Figs. 6c and 10c) that is in agreement with the maximum zero offset of the flow-meter transducers (see flow-meter characteristics in Section 2.1). The 0*k*[*rpm*] also highlights a modelling error in the simulation results, most clearly in the flow curves at Figs. 6a and 10a: the aortic flow wave produced by the pressure curve in the model is too triangular and short-timed, and the backflow during the valve closing is too large. Despite the poor overlap for the distributions seen in the ECDF plots Figs. 6d and 10d and the scatter plots Figs. 6c and 10c, the validation metrics in Figs. 6b and 10b for 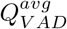 and 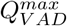 are almost negligible as the values enforced in the simulation are close to the experimental ones.

The 8*k*[*rpm*] operation condition provides the smallest overall metrics (i.e. best agreement) between the experimental and simulation. For the condition 22[%]@68.42[*bpm*] (Section 3.2.2) this can be seen as a good overlap between the experimental ranges and the simulation distributions. Similarly, the simulation ECDF curves (Fig. 7d) overlap the experiment and numerical ranges for every QoI. Almost similarly, the condition 17[%]@61.18[*bpm*] (Section 3.2.3) shows an overlap for most variables in the ECDF curves (Fig. 11d) except for those associated with the aortic flow (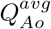 and 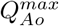). Despite that lack of overlap, the validation metrics in Fig. 11b are also reduced.

The lack of publications with UQ analyses for ventricular CFD makes comparing these results a difficult task. The results suggest a mismatch between the experimental and simulation results for the 0*k*[*rpm*] (LVAD off) and the 11*k*[*rpm*] cases. These validation points highlight potential issues that are worth exploring.

### 3.3 Discussion on the V&V40 credibility factors: achieved score

This section is intended to summarise the achieved scores for the credibility factors described in the ASME V&V40 [10] and summarised in Table 2. The section is intended to compare the pre-selected goal with the final achieved score. For the reader’s reference, the translation from the scores to the required activities can be found in the supporting material Section S1. The rationale behind the chosen goal for this work can be found in the supporting material Section S2. In the rest of this section we will summarise the rationale behind the scores obtained for each credibility factor.

#### Verification credibility factors

For the calculation and code verification the ASME standards define a set of rankings that ranges from no verification up to an extensive set of tests. The code used as simulation engine follows rigorous SQA enforced by the code life cycle tool (details on the supporting material Section S3.2.1). Also, the fact that the simulation code is partially open source and part of the Partnership for Advanced Computing in Europe (PRACE) Unified European Applications Benchmark Suite (UEABS) ensures the code transparency and constant scrutiny [48,49,50]. Added to this, two numerical code verification tests (supplementary material Section S3.2) and three numerical calculation verification tests (supplementary material Section S3.3) were executed to ensure code correctness and bounded numerical error for this problem. Comparing the executed tasks with the rankings in the supplementary material S1, led us to rank the SQA, NCV, discretisation error, and numerical solver error with the maximum score, surpassing the original desired goal. Due to the lack of external manpower, the inputs of the solver where only checked by internal review. This led us to achieve a C out of a maximum D in the user error credibility factor. As the number of input variables is rather small, the original goal was B out of D and therefore the original goal is achieved.

#### Validation credibility factors

While the background model is based the well known Navier-Stokes equations, there are multiple associated sub-models like the pump H-Q performance curve model, the lumped valve model or the Aortic impedance Windkessel model. Some of the assumptions for the simplification on the UQ were tested *a-priori* during the SA, therefore the model form correctness scores a B out of C, the desired goal. On the model input credibility factor, due to the thorough SA and UQ executed we achieved the maximum possible score, surpassing the original goals. The analysis was executed using retrospective experimental data, which was not gathered for VVUQ use. A single silicone ventricle was used as test sample, achieving an A out of C for the quantities of test samples credibility factor and an A out of D for the range of the test sample characteristics credibility factor. Despite this, the geometry used was characterised in all its features with a computer drawing tool, obtaining a score of C out of C. The CoU does not require evaluating the QoIs for different geometries, therefore a single idealised geometry was considered for the credibility evidence. With this, all the goals for the test sample credibility factors were achieved or surpassed. Further rationale can be found in the detailed table in the supplementary material S2. On the test conditions credibility factor, multiple test conditions were tested (achieving a B out of B) representing the expected extreme conditions range (achieving a C out of D), measuring all the key test conditions (achieving a B out of C). Despite this, the uncertainty of the test conditions was not characterised, achieving an A out of C. Most of the goals for the comparator credibility factor where achieved or surpassed, except for the characterisation of the test condition uncertainties. Finally in the assessment credibility factor, the accuracy of the simulation output is evaluated. As the simulation is designed to reproduce the experiment physics and resulting QoIs, all the input types and ranges of all inputs were identical,achieving a C out of C for the equivalence of inputs credibility factor. Also multiple outputs were rigorously compared via two approaches of validation metrics, achieving a C out of C for the equivalence, rigour and quantity of output variables credibility factor, surpassing the predefined goals in all of them. As the level of agreement was satisfactory for some key comparisons, we achieve the desired goal of B out of C for the agreement of output comparison credibility factor.

#### Applicability credibility factors

this item assesses how relevant the validation results are to the CoU. As the CoU proposes the tool for analysing LVAD and Aortic valve flow to analyse total cardiac output and aortic valve opening, we assign the maximum rank C out of C for the relevance of the QoIs for the question of interest. As the number of validation points could be increased to target a larger operation envelope, the validation activities only encompassed some validation points for the CoU, achieving a C out of B. In both cases, the goals where surpassed.

#### Overall credibility assessment

We demonstrate the model to be sufficiently close to the validation points for the simple CoU proposed. While some of the credibility factors did not obtained the maximum achievable score, they did obtained or surpassed the desired goal for the medium risk application (refer to Table 2 for a summary). Riskier applications, where the numerical model drive safety related conclusions or where the final decision relies more on middling, would require achieving those maximum achievable scores. Further improvements on the model to obtain these maximum scores are discussed in the conclusion.

## 4 Application to ramp study

As explained earlier, a ramp study [6] is routinely performed after LVAD implantation to select the pump speed for the patient. The selection is based on LV flow and geometry measured with echocardiography while the LVAD speed is increased over a wide range. The optimal speed is chosen to ensure end-organ perfusion, corresponding to a cardiac output (measured as 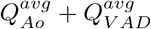) of 70[*cm*^3^/*s*] (assuming an average BSA of 1.9[*m*^2^]). Bowing of the intraventricular septum towards either RV (at low LVAD speeds) or LV (at high speeds) is avoided, particularly if the LVAD inflow cannula is oriented towards the AoV. It is desirable to achieve opening of the AoV (measured via a 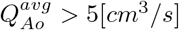), at least intermittently, which increases at lower LVAD speeds. The balance among these considerations are determined by the clinicians present, and the speed selected for long term LVAD support. Interestingly, most LVAD patients experience changes in cardiac geometry and function during the use of LVAD support. For example, increased heart rate or blood pressure can result in shorter systolic durations and alter AoV opening. Reduction in LV volume due to reverse remodeling and improvements in ejection fraction may shift the intra-ventricular septum position or enable greater AoV opening. With this, the final pump speed will depend on the patient’s hemodynamic condition, quantified here via the EF and HR. In all of these examples, having a more adaptive system begins with a validated tool such as the model proposed herein.

The model results extend the range of the experimental studies by evaluating the QoIs over a range of heart rate and ejection fraction, specifically AoV flow and total aortic flow (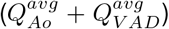). Figure 13 shows that the model predicts a small but notable decrease in LVAD flow with increasing HR and EF, which is accompanied by an increase in flow through the AoV. Figure 13 shows that the lower pump speed limit is bounded by 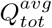 and the upper pump speed limit is bounded by 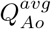. At 8*k*[*rpm*] the pump is unable to meet the 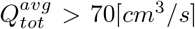 requirement for the range of HR and EF analysed. Oppositely, the 11*k*[*rpm*] case is unable to meet the 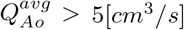 requirement for the situations with HR ≲ 65[*bpm*] or EF ≲ 18[%]. This agrees with the findings in [6] showing that the AoV closes at 9124 ± 1, 222[*rpm*] with an optimal LVAD speed of 8850 ± 470[*rpm*].

**Fig. 13:**
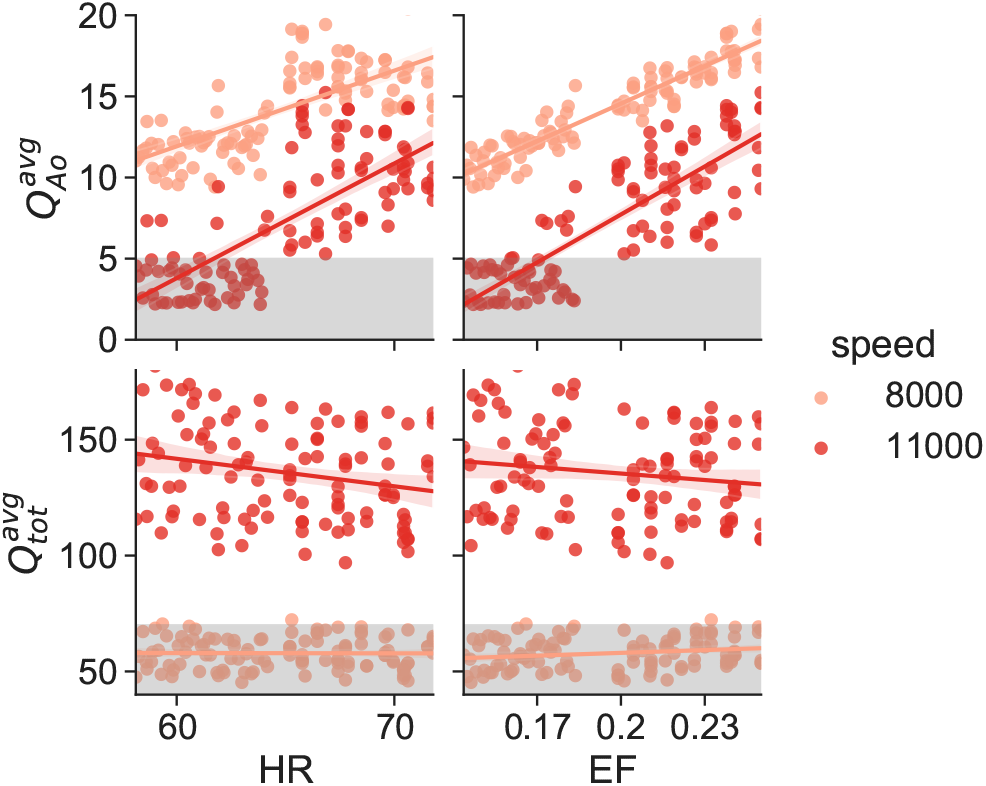
Aortic valve flow 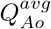 and total flow 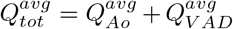 as a function of the HR and EF. Data is shown for both, 8*k*[*rpm*] and 11*k*[*rpm*]. The grey hatched region represents the minimum limit for both 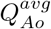 and 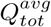.

Ideally, in future iterations of LVAD therapy, the device will be able to remotely monitor the HR, and the speed setting updated to adjust for the patient condition. With improvements in remote monitoring, device adjustment, the presented model could serve as a baseline tool for a smart interface between the patient’s LVAD and their clinical team, and form the foundation for automated speed control.

## 5 Conclusion

This manuscript provides a thorough detail and execution of a VVUQ plan of a clinically relevant numerical model. Starting from a major concern of LVAD treatment we define the V&V40 terminology and goals following the approved standard, and proceed to design a VVUQ plan that we afterwards execute and analyse with statistical tools.

While simpler LVAD numerical models have been published in the past, this is the first including a deformable ventricle, a pressure driven valve model and a dynamic LVAD boundary condition. The numerical model has been created to faithfully reproduce the SDSU-CS. To do so, the numerical model required to deform the mesh with the same pattern as the experiment, a 0D model of the systemic arteries, a novel approach for the valves that is driven by the transvalvular pressure gradient, and a novel approach to represent the LVAD through an H-Q curve performance function as boundary condition.

Moreover, such a model has been subject to the V&V40 pipeline allowing to bound the uncertainties in the simulation. The main facilitator for this has been the usage of a bench experiment as a source of comparators. While animal experimentation has a large inter-subject variability, low reproducibility and low access to the QoI, bench experiments provide a highly reproducible and accessible set of comparators. To ensure the solution procedure correctness, the numerical model was subjected to two code verification tests that bounded the numerical error for an operating condition close to the validation points. The two calculation verification tests executed, provided a measurement of the uncertainty produced by the spatial discretisation. The local SA provided a graphical understanding of the model’s behaviour, while the global SA based on total Sobol indices highlighted the most impactful input variables. This variable reduction brings the consequent reduction in the UQ analysis computational cost. The six validation points swiped through three pump speed velocities, two EF and two HR. While [51] reviews the use of computer model for critical health applications under the ASME V&V40 standard [10], this work presents for the first time a complete execution of a VVUQ plan following the cited guideline. The final use of the model is the application to the ramp study [6]. The model predicted an operational pump speed range that allows obtaining aortic valve opening and the desired total aortic flow. Moreover, the pump speed ranges predicted by the model agrees by the ranges found in [6].

Being the first iteration of the VVUQ plan, the results highlighted a number of items to be improved in the future. On the bench experiment side, including multiple executions for each validation point, would improve the validation metrics in the UQ, as the input variables could be characterised with a probability distribution instead of a forced 10% experimental uncertainty as done in the current manuscript. Retrieving multiple measurements of the H-Q curve of the LVAD for the operating condition will also have a direct positive impact on the final validation metrics, as we can conclude from the results in this work. The large uncertainty ranges in the H-Q curves in this manuscript produce distributions in the QoIs. On the simulation side, including valve geometries will improve the intra-LV flow patterns in future applications where vortex quantification becomes critical. Also, including the deterministic variables in the SA will increase the model credibility, necessary for a higher credibility score in the model form factor. These improvements will not only make a more accurate model, but also increase the scoring in the credibility factors. With these improvements, the model could target riskier applications.

## Acknowledgements

This project was funded in part by the FDA Critical Path Initiative and by an appointment to the Research Participation Program at the Division of Biomedical Physics, Office of Science and Engineering Laboratories, Center for Devices and Radiological Health, U.S. Food and Drug Administration, administered by the Oak Ridge Institute for Science, and Education through an interagency agreement between the U.S. Department of Energy and FDA. MV and AS acknowledge the funding from the projects `H2020-EU.1.4.1.3.` - CompBioMed2 - Grant agreement ID: `823712` and `H2020-EU.3.1.5.` - SilicoFCM - Grant agreement ID: `777204`. AS is partially funded by the Torres Quevedo Program (`PTQ2019-010528`). CB is partially funded by the Torres Quevedo Program (`PTQ2018-010290`). The authors want to extend their gratitude to Simone Venturi from Sandia National Laboratories and Tim Baldwin from the U.S. Food and Drug Administration (FDA) for the discussions about the clinical applications, about SA and UQ, for their reviews on the draft, and the learning material he provided. Finally, the authors want to acknowledge the crucial support of Adam J. Stephens from Sandia National Laborarories with the Dakota inputs.

## A Calculation of the pump input variable ranges

While the physical pump has the rotor speed as input, its operation is described through the pressure-flow curve, or simply the H-Q curve. The H-Q curve is a function of the pump speed and the characteristics of the inlet and outlet tubings, as these tubing affect the system pressure drop. To fit the pump H-Q curve model (Eq. (3) in Section 2.2.5), the SDSU recovered the H-Q performance curves of the used pump for multiple speeds. These measurements are shown as marks in Fig. 14. These measurements were fitted by a non-linear least squares method to a second order polynomial to obtain the LVAD pump coefficients (shown in Table 5) in Eq. (3). As, for this data *c*_VAD_ ~ 0.0 in every case, the quadratic coefficient was forced to zero *c*_VAD_ ≡ 0.0. The fittings are shown as lines in Fig. 14. The fitting error *ϵ_fit_*, shown as a light gray area in Fig. 14, is calculated as the square root of the diagonal of the fitting covariance matrix, also called the standard deviation. To account for experimental error *ϵ_exp_* we include a 10% error range for each coefficient, shown as dark gray in Fig. 14.

**Fig. 14:**
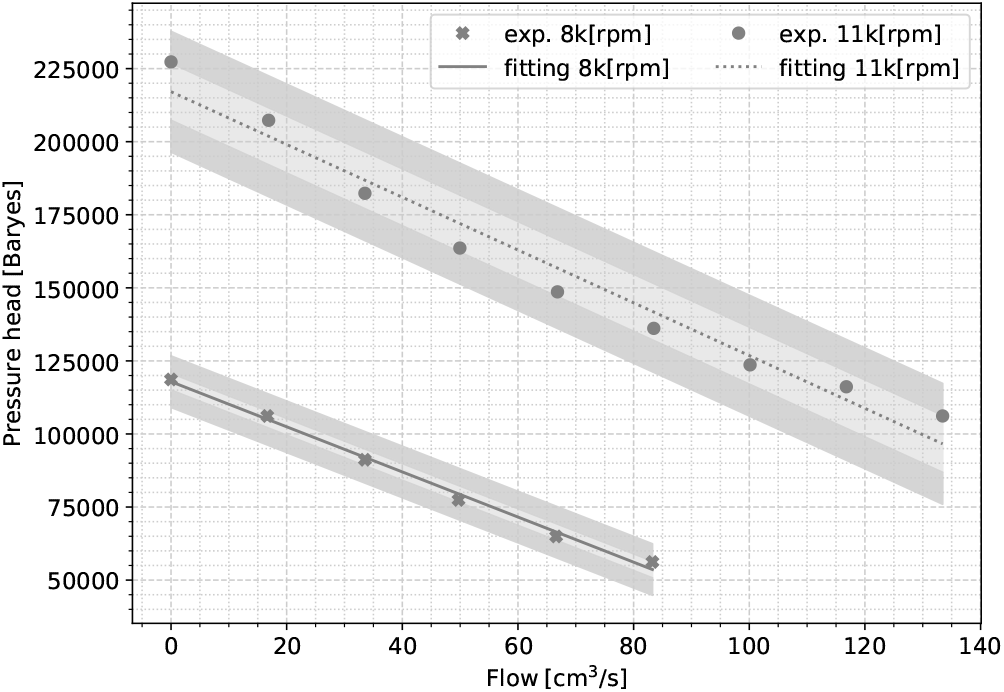
Different H-Q curves measured for the pump operating at multiple speeds (5*k*, 8*k*, 11*k*, 14*k*[*rpm*]). The light grey area is representing the measured fitting (numerical) error *ϵ_fit_* and the dark grey area is representing the assumed experimental error *ϵ_exp_*.

## Nomenclature

ALE: Arbitrarian Lagrangian-Eulerian. 1,5
AoV: Aortic Valve. 2, 19
ASGS: algebraic subgrid scale. 5
ASME: American Society of Mechanical Engineers. 3 7, 15, 16, 21
BSA: body surface area. 2, 19
BSC: Barcelona Supercomputing Center. 5
CDF: cumulative distribution function. 9
CFD: computational fluid dynamics. 3, 5, 9-11, 13, 15
CoU: context of use. 6-8, 17, 18
CS: cardiac simulator. 3, 9, 11, 21
DARE: Dakota server. 5, 6
ECDF: empirical cumulative distribution function. 9, 11 12,14-16, 18-20
EDV: end diastolic volume. 3-5
EF: Ejection Fraction. 4, 6, 10, 11, 19-21
ESV: end systolic volume. 4, 5
FDA: Food and Drug Administration. 3
FEM: finite elements method. 5
FSI: fluid-structure interaction. 3-5
GMRES: generalized minimal residual method. 5
HF: heart failure. 2, 4, 6
HPC: high performance computing. 6
HR: heart rate. 2, 6, 10-12, 19-21
KDE: kernel distribution estimation. 12, 14
LHS: latin hypercube sampling. 9, 10, 12
LV: left ventricle. 1-7, 9, 10, 18, 19, 21
LVAD: left ventricular assist device. 1-7, 9-12, 14-16, 18-22
MN: MinkowskiLļ norm. 9, 11-13
NCV: numerical code verification. 7, 16
NYHA: New York heart association. 4
PCE: polynomial chaos expansion. 9
PRACE: Partnership for Advanced Computing in Europe. 16
QoI: quantity of interest. 1, 6-15, 17-19, 21
RV: right ventricle. 2, 19
SA: sensitivity analysis. 5-11, 13, 16, 17, 21, 22
SDSU: San Diego State University. 3, 9, 11, 21, 22
SQA: software quality assurance. 7, 8, 16
UEABS: Unified European Applications Benchmark Suite. 16
UQ: uncertainty quantification. 4-13, 15-17, 21, 22
VVUQ: verification, validation and uncertainty quantification. 1,3, 6, 7, 9, 17, 21

## Footnotes

1 resolution: 50[*ml/min*], max zero offset: 0.3[*ml/min*], absolute accuracy: 10%

2 resolution: 10[*ml/min*], max zero offset: 0.06[*ml/min*], absolute accuracy: 10%.

3 sensitivity: 5 ± 1%[*mmHg*], zero offset: 25[*mmHg*].

4 sampling frequency: 200[*Hz*], impedance 1[*M*Ω]@1[*pF*], resolution: 16[*bit*].

5 version: 1.8.7.

## References

[1] Benjamin, E. J., Muntner, P., Alonso, A., Bittencourt, M. S., Callaway, C. W., Carson, A. P., Chamberlain, A. M., Chang, A. R., Cheng, S., Das, S. R., et al., 2019. “Heart disease and stroke statistics—2019 update: a report from the american heart association”. Circulation, 139(10), pp. e56–e528.

[2] Fang, J. C., Ewald, G. A., Allen, L. A., Butler, J., Canary, C. A. W., Colvin-Adams, M., Dickinson, M. G., Levy, P., Stough, W. G., Sweitzer, N. K., et al., 2015. “Advanced (stage d) heart failure: a statement from the heart failure society of america guidelines committee”. Journal of cardiac failure, 21(6), pp. 519–534.

[3] Everly, M. J., 2008. “Cardiac transplantation in the united states: an analysis of the unos registry.”. Clinical transplants, pp. 35–43.

[4] Jorde, U. P., Kushwaha, S. S., Tatooles, A. J., Naka, Y., Bhat, G., Long, J. W., Horstmanshof, D. A., Kormos, R. L., Teuteberg, J. J., Slaughter, M. S., et al., 2014. “Results of the destination therapy post-food and drug administration approval study with a continuous flow left ventricular assist device: a prospective study using the intermacs registry (interagency registry for mechanically assisted circulatory support)”. Journal of the American College of Cardiology, 63(17), pp. 1751–1757.

[5] Topilsky, Y., Hasin, T., Oh, J. K., Borgeson, D. D., Boilson, B. A., Schirger, J. A., Clavell, A. L., Frantz, R. P., Tsutsui, R., Liu, M., et al., 2011. “Echocardiographic variables after left ventricular assist device implantation associated with adverse outcome”. Circulation: Cardiovascular Imaging, 4(6), pp. 648–661.

[6] Uriel, N., Morrison, K. A., Garan, A. R., Kato, T. S., Yuzefpolskaya, M., Latif, F., Restaino, S. W., Mancini, D. M., Flannery, M., Takayama, H., et al., 2012. “Development of a novel echocardiography ramp test for speed optimization and diagnosis of device thrombosis in continuous-flow left ventricular assist devices: the columbia ramp study”. Journal of the American College of Cardiology, 60(18), pp. 1764–1775.

[7] Jorde, U. P., Uriel, N., Nahumi, N., Bejar, D., Gonzalez-Costello, J., Thomas, S. S., Han, J., Morrison, K. A., Jones, S., Kodali, S., et al., 2014. “Prevalence, significance, and management of aortic insufficiency in continuous flow left ventricular assist device recipients”. Circulation: Heart Failure, 7(2), pp. 310–319.

[8] Roy, C. J., and Oberkampf, W. L., 2011. “A comprehensive framework for verification, validation, and uncertainty quantification in scientific computing”. Computer methods in applied mechanics and engineering, 200(25-28), pp. 2131–2144.

[9] Test Uncertainty, A., 2006. “Ptc 19.1-2005”. American Society of Mechanical Engineers, 3, pp. 10016–5990.

[10] American Society of Mechanical Engineers, 2018. “Assessing Credibility of Computational Modeling through Verification and Validation: Application to Medical Devices - V V 40 - 2018”. Asme V&V 40-2018, p. 60.

[11] American Society of Mechanical Engineers, 2009. “Standard for Verification and Validation in Computational Fluid Dynamics and Heat Transfer: ASME V&V 20”. The American Society of Mechanical Engineers (ASME).

[12] Chivukula, V. K., Beckman, J. A., Prisco, A. R., Lin, S., Dardas, T. F., Cheng, R. K., Farris, S. D., Smith, J. W., Mokadam, N. A., Mahr, C., et al., 2019. “Small lv size is an independent risk factor for vad thrombosis”. ASAIO journal (American Society for Artificial Internal Organs: 1992), 65(2), p. 152.

[13] Prisco, A. R., Aliseda, A., Beckman, J. A., Mokadam, N. A., Mahr, C., and Garcia, G. J., 2017. “Impact of lvad implantation site on ventricular blood stagnation”. ASAIO journal (American Society for Artificial Internal Organs: 1992), 63(4), p. 392.

[14] Ong, C., Dokos, S., Chan, B., Lim, E., Al Abed, A., Osman, N., Kadiman, S., and Lovell, N. H., 2013. “Numerical investigation of the effect of cannula placement on thrombosis”. Theoretical Biology and Medical Modelling, 10(1), pp. 1–14.

[15] Liao, S., Neidlin, M., Li, Z., Simpson, B., and Gregory, S. D., 2018. “Ventricular flow dynamics with varying lvad inflow cannula lengths: In-silico evaluation in a multiscale model”. Journal of biomechanics, 72, pp. 106–115.

[16] Chivukula, V. K., Beckman, J. A., Li, S., Masri, S. C., Levy, W. C., Lin, S., Cheng, R. K., Farris, S. D., Wood, G., Dardas, T. F., et al., 2020. “Left ventricular assist device inflow cannula insertion depth influences thrombosis risk”. Asaio Journal, 66(7), pp. 766–773.

[17] Neidlin, M., Liao, S., Li, Z., Simpson, B., Kaye, D. M., Steinseifer, U., and Gregory, S., 2021. “Understanding the influence of left ventricular assist device inflow cannula alignment and the risk of intraventricular thrombosis”. Biomedical engineering online, 20(1), pp. 1–14.

[18] Wong, K., Samaroo, G., Ling, I., Dembitsky, W., Adamson, R., Del Álamo, J., and May-Newman, K., 2014. “Intraventricular flow patterns and stasis in the lvad-assisted heart”. Journal of biomechanics, 47(6), pp. 1485–1494.

[19] Garcia, M. A. Z., Enriquez, L. A., Dembitsky, W., and May-Newman, K., 2008. “The effect of aortic valve incompetence on the hemodynamics of a continuous flow ventricular assist device in a mock circulation”. ASAIO journal, 54(3), pp. 237–244.

[20] Segur, J. B., and Oberstar, H. E., 1951. “Viscosity of glycerol and its aqueous solutions”. Industrial & Engineering Chemistry, 43(9), pp. 2117–2120.

[21] Yin, F., and Liu, Z., 1989. “Estimating arterial resistance and compliance during transient conditions in humans”. American Journal of Physiology-Heart and Circulatory Physiology, 257(1), pp. H190–H197.

[22] May-Newman, K., Enriquez-Almaguer, L., Posuwattanakul, P., and Dembitsky, W., 2010. “Biomechanics of the aortic valve in the continuous flow vad-assisted heart”. Asaio Journal, 56(4), pp. 301–308.

[23] Association, N. Y. H., 1964. “Diseases of the heart and blood vessels: nomenclature and criteria for diagnosis”. Little, Brown.

[24] Santiago, A., Zavala-Aké, M., Borell, R., Houzeaux, G., and Vázquez, M., 2020. “Hpc compact quasi-newton algorithm for interface problems”. Journal of Fluids and Structures, 96, p. 103009.

[25] Vázquez, M., Houzeaux, G., Koric, S., Artigues, A., Aguado-Sierra, J., Arís, R., Mira, D., Calmet, H., Cucchietti, F., Owen, H., et al., 2016. “Alya: Multiphysics engineering simulation toward exascale”. Journal of computational science, 14, pp. 15–27.

[26] Calderer, R., and Masud, A., 2010. “A multiscale stabilized ALE formulation for incompressible flows with moving boundaries”. Computational Mechanics, 46(1), pp. 185–197.

[27] Codina, R., 2000. “Stabilization of incompressibility and convection through orthogonal subscales in finite element methods”. Computer Methods in Applied Mechanics and Engineering, 190(13-14), pp. 1579–1599.

[28] Houzeaux, G., Vázquez, M., Aubry, R., and Cela, J. M., 2009. “A massively parallel fractional step solver for incompressible flows”. Journal of Computational Physics, 228(17), pp. 6316–6332.

[29] Houzeaux, G., Aubry, R., and M. Vazquez, 2011. “Extension of fractional step techniques for incompressible flows: The preconditioned orthomin(1) for the pressure schur complement”. Computers & Fluids, 44(1), pp. 297–313.

[30] Santiago, A., Aguado-Sierra, J., Zavala-Aké, M., Doste-Beltran, R., Gómez, S., Arís, R., Cajas, J. C., Casoni, E., and Vázquez, M., 2018. “Fully coupled fluid-electro-mechanical model of the human heart for supercomputers”. International journal for numerical methods in biomedical engineering, 34(12), p. e3140.

[31] BM, A., J, W., KR, D., JP, E., and MS, E., 2009. “Dakota, a multilevel parallel object-oriented framework for design optimization, parameter estimation, uncertainty quantification, and sensitivity analysis: version 5.0 user’s manual”. Sandia National Laboratories, Tech. Rep. SAND2010-2183.

[32] B.M., A., Bohnhoff, W., K.R., D., M.S., E., J.P., E., M.S., E., G., G., R.W., H., Hough, P., and K.T., H., 2019. White paper: Programming according to the fences and gates model for developing assured, secure software systems. Tech. Rep. SAND2014-4633, Dakota, A Multilevel Parallel Object-Oriented Framework for Design Optimization, Parameter Estimation, Uncertainty Quantification, and Sensitivity Analysis: Version 6.11 User’s Manual, November.

[33] Morrison, T. M., Hariharan, P., Funkhouser, C. M., Afshari, P., Goodin, M., and Horner, M., 2019. “Assessing Computational Model Credibility Using a Risk-Based Framework”. ASAIO Journal.

[34] Trucano, T. G., Swiler, L. P., Igusa, T., Oberkampf, W. L., and Pilch, M., 2006. “Calibration, validation, and sensitivity analysis: What’s what”. Reliability Engineering & System Safety, 91(10-11), pp. 1331–1357.

[35] Pearson, K., 1895. “Vii. note on regression and inheritance in the case of two parents”. proceedings of the royal society of London, 58(347-352), pp. 240–242.

[36] Sobol, I. M., 2001. “Global sensitivity indices for nonlinear mathematical models and their monte carlo estimates”. Mathematics and computers in simulation, 55(1-3), pp. 271–280.

[37] Najm, H. N., 2009. “Uncertainty quantification and polynomial chaos techniques in computational fluid dynamics”. Annual review of fluid mechanics, 41, pp. 35–52.

[38] Voyles, I. T., and Roy, C. J., 2015. “Evaluation of model validation techniques in the presence of aleatory and epistemic input uncertainties”. In 17th AIAA Non-Deterministic Approaches Conference, p. 1374.

[39] Levrero-Florencio, F., Margara, F., Zacur, E., Bueno-Orovio, A., Wang, Z., Santiago, A., Aguado-Sierra, J., Houzeaux, G., Grau, V., Kay, D., et al., 2020. “Sensitivity analysis of a strongly-coupled human-based electromechanical cardiac model: Effect of mechanical parameters on physiologically relevant biomarkers”. Computer methods in applied mechanics and engineering, 361, p. 112762.

[40] Aguado-Sierra, J., Butakoff, C., Brigham, R., Baron, A., Houzeaux, G., Guerra, J. M., Carreras, F., Filgueiras-Rama, D., Iaizzo, P. A., Iles, T. L., et al., 2021. “In-silico clinical trial using high performance computational modeling of a virtual human cardiac population to assess drug-induced arrhythmic risk”. medRxiv.

[41] Pathmanathan, P., Cordeiro, J. M., and Gray, R. A., 2019. “Comprehensive uncertainty quantification and sensitivity analysis for cardiac action potential models”. Frontiers in physiology, 10, p. 721.

[42] Tolpen, S., Janmaat, J., Reider, C., Kallel, F., Farrar, D., and May-Newman, K., 2015. “Programmed speed reduction enables aortic valve opening and increased pulsatility in the lvad-assisted heart”. Asaio Journal, 61(5), pp. 540–547.

[43] Mudd, J. O., Cuda, J. D., Halushka, M., Soderlund, K. A., Conte, J. V., and Russell, S. D., 2008. “Fusion of aortic valve commissures in patients supported by a continuous axial flow left ventricular assist device”. The Journal of heart and lung transplantation, 27(12), pp. 1269–1274.

[44] Gewillig, M., Brown, S. C., Eyskens, B., Heying, R., Ganame, J., Budts, W., Gerche, A. L., and Gorenflo, M., 2010. “The fontan circulation: who controls cardiac output?”. Interactive cardiovascular and thoracic surgery, 10(3), pp. 428–433.

[45] Vincent, J.-L., 2008. “Understanding cardiac output”. Critical care, 12(4), pp. 1–3.

[46] Saks, V., Dzeja, P., Schlattner, U., Vendelin, M., Terzic, A., and Wallimann, T., 2006. “Cardiac system bioenergetics: metabolic basis of the frank-starling law”. The Journal of physiology, 571(2), pp. 253–273.

[47] Campos, J., Sundnes, J., Dos Santos, R., and Rocha, B., 2020. “Uncertainty quantification and sensitivity analysis of left ventricular function during the full cardiac cycle”. Philosophical Transactions of the Royal Society A, 378(2173), p. 20190381.

[48] Bulla, J. M., and Emerson, A., 2019. Selection of a unified european application benchmark suite. Tech. rep., Partnership for Advanced Computing in Europe (PRACE).

[49] Rodriguez, J., 2019. Performance analysis of alya on a tier-0 machine using extrae. Tech. rep., Partnership for Advanced Computing in Europe (PRACE).

[50] Houzeaux, G., and Artigues, T., 2016. Parallel mesh partitioning in alya. Tech. rep., Partnership for Advanced Computing in Europe (PRACE).

[51] Parvinian, B., Pathmanathan, P., Daluwatte, C., Yaghouby, F., Gray, R. A., Weininger, S., Morrison, T. M., and Scully, C. G., 2019. “Credibility evidence for computational patient models used in the development of physiological closed-loop controlled devices for critical care medicine”. Frontiers in physiology, 10, p. 220.

