## Supplementary Material for "Design and execution of a Verification, Validation, and Uncertainty Quantification plan for a numerical model of left ventricular flow after LVAD implantation"

### Supporting Material


Alfonso Santiago[1,2], Constantine Butakoff[2], Beatriz Eguzkitza[1], Richard A. Gray[3], Karen May-Newman[4], Pras Pathmanathan[3], Vi Vu[4], Mariano Vázquez[1,2]

[1] Barcelona Supercomputing Center (BSC), Barcelona, 08034, Spain. [2] ELEM biotech, Barcelona, 08034, Spain.. [3] US Food and Drug Administration (FDA), Silver Spring (MD), USA.. [4] Department of Mechanical Engineering, San Diego State University (SDSU) San Diego (CA), USA..


#### Nomenclature

**BDF** backward differentiation formula. 7, 8
**CD/CI** continuous integration and continuous deployment. 6
**CFD** computational fluid dynamics. 8, 9
**CoU** context of use. 4, 5
**FEM** finite elements method. 6
**GCI** grid convergence index. 6
**LV** left ventricle. 9
**LVAD** left ventricular assist device. 8
**MMS** method of manufactured solutions. 4
**QoI** quantity of interest. 2–6
**RMSE** root mean square error. 6, 8, 9
**SA** sensitivity analysis. 4
**SQA** software quality assurance. 1, 4
**UQ** uncertainty quantification. 5

### S1 Ranking for risk informed credibility assessment

This section provides the gradiation for each credibility factor and actions required on each item in the standard ASME V&V40 [1].

1. **VERIFICATION:** Activities related to the correctness of the implementation of the numerical model.
   1.1. **Code verification:** Activities to ensure the correct implementation of the code.
      1.1.1. **Software quality assurance (SQA):** Activities to ensure repeatability and traceability of the code modifications. Ranked in three:
         A. Very little or no software quality assurance (SQA) are followed.
         B. SQA procedures are specified and documented.
         C. SQA procedures are specified and documented. Quality metrics are tracked. Code anomalies are systematically registered and tracked.
      1.1.2. **Numerical code verification (NCV)**: Activities related to demonstrate the correct implementation and functioning of the numerical algorithms. Ranked in four:
         A. No NCV was performed.
         B. The numerical solution is compared to a benchmark solution from another code.
         C. The numerical solution is compared to an exact analytical or manufactured solution, demonstrating an asymptotical approach with mesh size.
         D. The order of accuracy is compared to the theoretical order of accuracy in an exact solution.

1.2. **Calculation verification:** Estimate the numerical error in the quantity of interests (QoIs) due to spatial and temporal discretisation.

1.2.1. **Discretisation error:** Estimation of error due to the finite points in time/space in which the problem is solved. Ranked in three:

A. No space/time convergence analysis is performed.
B. Space and time convergence analysis are performed obtaining stable behaviours.
C. Grid and space convergence analysis is performed, estimating the discretisation error.

1.2.2. **Numerical solver error:** Errors originated from the numerical solution based on the solver parameters.

A. No solver parameter sensitivity was performed.
B. Solver parameters are based on values from a previously verified model.
C. A solver parameter sensitivity study is performed ensuring that the chosen values does have a negligible impact in the final model accuracy.

1.2.3. **User error:** refers to errors accrued by the practitioner (unchecked inputs).

A. Inputs and outputs were not verified.
B. Key inputs and outputs were verified by the practitioner.
C. Key inputs and outputs were verified by an internal peer review.
D. Key inputs and outputs were verified by reproducing simulations by an external reviewer.

2. **VALIDATION:** Process of assessing the degree to which the computational model is an appropriate representation of the reality for the context of use.

2.1. **Computational model:** Refers to the input of the numerical model.

2.1.1. **Model form:** Refers to the correctness of the conceptual and mathematical formulation of the computational model. Ranked on three:

A. Influence of the model assumptions are not explored.
B. Influence of some assumptions is explored.
C. Influence of every assumption is explored.

2.1.2. **Model inputs:** Refer to the values of parameters used.

2.1.2.1. **Quantification of sensitivities:** examines the degree to which the model's output is sensitive to the model inputs. Ranked on three:

A. A sensitivity analysis is not performed.
B. A sensitivity analysis of the expected key parameters is performed.
C. Comprehensive sensitivity analysis is performed.

2.1.2.2. **Quantification of uncertainties:** the degree to which known or assumed uncertainties in the model inputs are propagated to uncertainties in the simulation. Ranked in four:

A. Uncertainties are not quantified.
B. Uncertainties on expected key inputs are identified and quantified but not propagated to assess the effect in the QoIs.
C. Uncertainties on expected key inputs are identified, quantified and propagated to assess the effect in the QoIs.
D. Uncertainties on all inputs are identified and quantified and propagated to assess the effect of the simulation results.

2.2. **Comparator**: Is the data against which the simulation results are evaluated.

2.2.1. **Test samples:** Refers to the population and characteristics of the experimental subjects.

2.2.1.1. **Quantity of test samples:** examines the number of samples used. Ranked in three:

A. A single sample is used.
B. Multiple samples are used, but not being statistically relevant.
C. A statistically relevant number of samples are used.

2.2.1.2. **Range of characteristics of test samples:** This item examines the number of test conditions used. Ranked in four:

A. A single test condition is examined.
B. Test conditions in a nominal range are examined.
C. Extreme test conditions are examined.
D. The entire range of test conditions is examined.

2.2.1.3. **Measurements of test samples:** Evaluate the rigor with which the measurement data characterize each test sample. Ranked in three:

A. The test sample is not characterized (measured).
B. One or more key characteristic are measured.
C. All key characteristics are measured.

2.2.1.4. **Uncertainty of test samples measurements:** This factor examines the analysis of the uncertainty associated with the tools and methods used. Ranked in four:

A. Characteristics uncertainty not addressed.
B. Uncertainty analysis incorporates instrument accuracy only.
C. Uncertainty analysis incorporates instrument accuracy and statistics (repeated measurements).

2.2.2. **Test conditions:** evaluate the rigorousness in which the tests were executed.

2.2.2.1. **Quantity of test conditions:** Number of test conditions imposed and characterized. Ranked in two:

A. Single test condition.
B. Multiple test conditions.

2.2.2.2. **Range of test conditions:** evaluates the range of test conditions included in the comparator study. Ranked in four:

A. A single test condition is examined.
B. Test conditions representing a range of conditions near nominal range are examined.
C. Test conditions representing the expected extreme conditions are examined.
D. Test conditions representing the entire range of conditions is examined.

2.2.2.3. **Measurements of test conditions:** Examines the rigor with the measurement data that characterize the test conditions. Ranked in three:

A. The test conditions are not measured.
B. One or more key test conditions are measured.
C. All key test conditions are measured.

2.2.2.4. **Uncertainty of test conditions:** This component analyses the uncertainty associated with the tools and methods to characterize the test conditions. Ranked in four:

A. Test conditions were not characterized or their uncertainty analysis is not executed.
B. Uncertainty analysis of the test conditions characteristics incorporated instrument accuracy only.
C. Uncertainty analysis of the test conditions characteristics incorporate instrument accuracy and statistics (repeated measurements).

2.3. **Assessment:** of the accuracy of the simulation output.

2.3.1. **Equivalence of input parameters:** between the type and range of input parameters. Ranked in three:

A. The types of some inputs are dissimilar.
B. The types of all inputs are similar, but ranges were not equivalent.
C. The types and ranges of all inputs are similar.

2.3.2. **Output comparison:** Equivalency between the types of output from the computational model and those from the comparator leads to increased credibility.

2.3.2.1. **Quantity:** Quantity of QoIs to compare. Ranked in two:

A. A single output is compared.
B. Multiple outputs are compared.

2.3.2.2. **Equivalence of output parameters:** Referring to the types of outputs to be compared. Ranked in three:

A. Most types of outputs are dissimilar.
B. Most types of outputs are similar.
C. Most types of outputs are equivalent.

2.3.2.3. **Rigor of output comparison:** This refers to the method used to compare the QoIs from the computational model:

A. Visual comparison is performed.
B. Comparison is performed by arithmetic difference.
C. Uncertainty in the output of the computational model or the comparator was used.

2.3.2.4. **Agreement of output comparison:** qualitative or quantitative agreement between the QoIs in the computational model and the comparator:

A. The level of agreement is not satisfactory for key comparison.
B. The level of agreement is satisfactory for some key comparisons.
C. The level of agreement is satisfactory for all comparisons.

3. **APPLICABILITY:** Attains the relevance of the validation to support the use of the model for a determined Context of Use.

3.1. **Relevance of the Quantities of Interest for the Question of Interest**: this compares the QoIs from the validation activities to the QoIs for the context of use (CoU). Ranked in three:

A. The QoIs from validation are related but not identical to those for the CoU.
B. A subset of the QoIs from the validation are identical to those for the CoU.
C. The QoIs from the validation are identical to those for the CoU.

3.2. **Relevance of the validation activities to the CoU:** This factor summarizes the relative proximity of the CoU to the validation points. Ranked in four:

A. There was no overlap between the ranges of the validation points and the CoU.
B. There was partial overlap between the ranges of the validation points and the CoU.
C. The CoU encompassed some validation points.
D. The CoU encompassed all validation points.

### S2 Rationale behind the achieved scores for each credibility factor

This section provides the rationale behind every achieved score for each credibility factor. The score is taken frm Section S1.

1. **VERIFICATION:**

1.1. **Code verification:**

1.1.1. **Software quality assurance (SQA).** *Maximum ranking: (C). Selected goal: (B). Achieved: (C).* The simulation engine used is an industrial code in constant evolution and therefore every issue and feature is tracked and metrics are periodically computed to ensure repeatably of the numerical results. The developers and expert users have the option to access the documentation and a SQA platform to report abnormal behaviour.

1.1.2. **Numerical code verification (NCV).** *Maximum ranking: (D). Selected goal: (C). Achieved: (D).* Predictions are based on the correct implementation solution of the incompressible Navier-Stokes model. Multiple cases of the method of manufactured solutions (MMS) were used for a grid convergence study and to evaluate the observed order of accuracy.

1.2. **Calculation verification:**

1.2.1. **Discretisation error** *Maximum ranking: (C). Selected goal (B). Achieved: (C).* The simulation involves the computation of QoIs that might be sensible to spatial discretisation. Space and time convergence analysis were performed estimating the discretisation error for the problem-specific QoIs.

1.2.2. **Numerical solver error.** *Maximum ranking: (C). Selected goal: (B). Achieved: (B).* The main QoI, this is the velocity field, is robust to solver parameters. Therefore solver parameters are based on values form previous executions.

1.2.3. **User error.** *Maximum ranking: (D). Selected goal: (B). Achieved: (C).* The number of input physical parameters is rather small ($\sim 5$), leading to little chance on having user error. Despite this, all inputs were verified by an internal peer review.

2. **VALIDATION:**

2.1. **Computational model:**

2.1.1. **Model form.** *Maximum ranking: (C). Selected goal: (B). Achieved: (C).* The complex model involves multiple assumptions. Despite this, it's based in known and proven equations.

2.1.2. **Model inputs:**

2.1.2.1. **Quantification of sensitivities.** *Maximum ranking: (C). Selected goal: (B). Achieved: (C).* Despite being a relatively reduced number of input variables the model is complex. A comprehensive sensitivity analysis (SA) was executed for the input variables. The variables classified as deterministic will be associated with a rationale that justify such classification.

2.1.2.2. **Quantification of uncertainties.** *Maximum ranking: (D). Selected goal: (B). Achieved: (D).* The background experiment used for the validation is highly controlled and reproducible, therefore capable of obtaining statistical measures for the input variables and QoIs. A comprehensive uncertainty quantification (UQ) was executed for the combined epistemic-aleatory variables, propagating the effect of their uncertainties.

2.2. **Comparator**:

2.2.1. **Test samples:**

2.2.1.1. **Quantity of test samples.** *Maximum ranking: (C). Selected goal: (A). Achieved: (A).* The silicone ventricle used in the bench experiment is CAD-designed and manufactured by casting. Therefore, reproducibility of the geometry is ensured and no more than one silicone ventricle sample is required.

2.2.1.2. **Range of characteristics of test samples.** *Maximum ranking: (D). Selected goal: (A). Achieved: (A).* A single test sample is used in the nominal range of key characteristics.

2.2.1.3. **Measurements of test samples.** *Maximum ranking: (C). Selected goal: (C). Achieved: (C).* The bench experiment is designed to be reproducible and in full control by the experimentalist. The ventricle is produced from a computer draw and therefore all the key characteristics of the sample are measured and easily reproduced

2.2.1.4. **Uncertainty of test samples measurements.** *Maximum ranking: (C). Selected goal: (A). Achieved: (B).* As a single test sample was used, uncertainty of characteristics is not quantified nor required.

2.2.2. **Test conditions:**

2.2.2.1. **Quantity of test conditions.** *Maximum ranking: (B). Selected goal: (B). Achieved: (B).* To ensure predictability of the computational model, multiple test conditions are evaluated.

2.2.2.2. **Range of test conditions.** *Maximum ranking: (D). Selected goal: (B). Achieved: (C).* The computational model was validated in extreme conditions for the pump speed.

2.2.2.3. **Measurements of test conditions.** *Maximum ranking: (C). Selected goal: (B). Achieved: (B).* The easy access for measurements of the bench experiment allows measuring multiple test conditions.

2.2.2.4. **Uncertainty of test conditions.** *Maximum ranking: (C). Selected goal: (B) Achieved: (A).* As the work is executed with retrospective bench data, the test conditions are not characterised nor their uncertainty analysed.

2.3. **Assessment:**

2.3.1. **Equivalence of input parameters.** *Maximum ranking: (C). Selected goal: (B). Achieved: (C).* As the model deals with classical fluid dynamics where experimental parameters are easy to measure and implement in computational model. Therefore all types and ranges of all inputs were similar.

2.3.2. **Output comparison:**

2.3.2.1. **Quantity.** *Maximum ranking: (B). Selected goal: (B). Achieved: (B).* Multiple QoIs extracted from the flow meters are compared.

2.3.2.2. **Equivalence of output parameters.** *Maximum ranking: (C). Selected goal: (B). Achieved: (C).* The goal of the current validation plan is to reproduce exactly the same measurements in the numerical model as in the experimental benchmark, therefore all the outputs are equivalent.

2.3.2.3. **Rigor of output comparison.** *Maximum ranking: (C). Selected goal: (B). Achieved: (C).* The outputs were compared with multiple validation metrics, therefore a rigorous comparison was executed.

2.3.2.4. **Agreement of output comparison.** *Maximum ranking: (CD). Selected goal: (B). Achieved: (C).* Most of the characteristic had satisfactory agreement but some of them only partially agreed.

3. **APPLICABILITY:**

3.1. **Relevance of the QoIs for the Question of interest** *Maximum ranking: (C). Selected goal: (B). Achieved: (C).* The numerical model is designed to retrieve the same QoIs as the required for the Question of interest, and these QoIs quantified during the UQ. Therefore the QoIs from the validation were identical to those for the CoU.

3.2. **Relevance of the validation activities to the CoU** *Maximum ranking: (D). Selected goal: (B). Achieved: (C).* The number of validation points were scarce due to limited available retrospective data.

### S3 Verification

#### S3.1 Mesh convergence metrics

The root mean square error (RMSE) between the solution $i$ and $j$ is defined thought the L2 norm $||.||_2$ as:

$$\epsilon_{i,j} = ||v_i - v_j|| = \sqrt{(\tilde{v}_i - \tilde{v}_j)^2} \quad (1)$$

Once all the cases are computed and the errors $\epsilon_{1,2}$ and $\epsilon_{2,3}$ computed the observed order of convergence is calculated as [2]:

$$s_{i,k} = 1 \cdot sign\left(\epsilon_{i,j}/\epsilon_{i,j}\right) \quad (2)$$

$$q_{i,k}(p_{i,k}) = \ln\left(\frac{r_{j,k}^{p_{i,k}} - s}{r_{i,j}^{p_{i,k}} - s}\right) \quad (3)$$

$$p_{i,k} = \left[1/\ln(r_{j,k})\right]\left[\ln|\epsilon_{i,j}/\epsilon_{j,k}| + q_{i,k}(p_{i,k})\right] \quad (4)$$

Note that in our case $r_{i,j} = r_{j,k} = r = 2.0$, so $q_{i,k}(p_{i,k}) = 0.0$ and therefore the previous system of equations is reduced to:

$$p_{i,k} = \frac{\ln|\epsilon_{i,j}/\epsilon_{j,k}|}{\ln(r)} \quad (5)$$

For three velocity fields computed $u_1, u_2, u_3$, $u_1$ being the coarsest, for three subdivision levels, the order of the convergence of the numerical scheme is given by:

$$p = \frac{\ln(||u_1 - u_2||_2/||u_2 - u_3||_2)}{\ln(2.0)} \quad (6)$$

With the observed $p$ value, the grid convergence index (GCI) can be computed as [2]:

$$GCI_{i,j}^{95\%} = 1.25\frac{\epsilon_{i,j}}{r_{i,j}^{p} - 1} \quad (7)$$

This uncertainty estimate provides an interval $f \pm U^{95\%}$ within the true mathematical value $f_T$ falls with a probability of $95\%$.

#### S3.2 Numerical code verification

Numerical code verification is executed as per Section 2 of [3], for a 2D Poiseuille and a 3D Womersley flow problem in a cylindrical tube. These problems have non-trivial analytical solutions that are used as true value. For both cases the discretisation error is monitored as the grid is systematically refined by halving as in [4]. If the ratio between mesh subdivisions is defined as $r_{i,j} = r_i/r_j$ then, for this case $r_{1,2} = r_{2,3} = r = 2.0$, a figure considerably larger than 1.3, the minimum value recommended [3]. The velocity field is the QoI to be verified as it is also the raw variable used calculated in the numerical model. The mesh convergence metrics are described in Section S3.1.

##### S3.2.1 Software Quality Assurance

The finite elements method (FEM) software is developed with a continuous integration and continuous deployment (CD/CI) strategy based on Git, which combines feature-driven development and feature branches with issue tracking. Git pipelines ensure continuous integration, running a series of software checks, builds, and regression tests when the developers modify the source code. The build evaluation includes 27 combinations of architectures (Intel, IBM), compilers (gnu, intel, pg, xl), and optimization options, running more than 200 regression tests executed with various MPI and OpenMP configurations, more than 4000 different executions. This method helps

Table 1: RMSE and RMSE expressed as percentage of 1.25[cm/s] for the Poiseuille flow problem solution in 2D.

| halving $[-]$ | Elements $[-]$ | Time step $[s]$ | RMSE $[cm/s]$ | RMSE [%] | $GCI^{95\%}$ $[cm/s]$ |
|---|---|---|---|---|---|
| 0 | 12,800 | 0.01 | 0.004555 | 0.36 | 0.00206 |
| 1 | 51,200 | 0.005 | 0.001361 | 0.11 | 0.000617 |
| 2 | 204,800 | 0.0025 | 0.000379 | 0.03 | 0.000171 |

to detect bugs early in the development cycle and guarantee the correctness of the simulation results and the software's stability. On this manuscript Alya version `v1.0-4454-gf0306e2d8` was used.

##### S3.2.2 2D Poiseuille

**Problem description:** This test involves a constant Poiseuille flow in a 2D rectangle.

**Domain definition and space-time discretisation:** The mesh is a rectangle $0.8 \times 4[cm]$, centered at and aligned with $X$ axis. Three meshes with 12800, 51200 and, 204800 elements were used (ratio $r_{i,j} = 2.0$). For each mesh a time step of 0.01$[s]$, 0.005$[s]$ and, 0.0025$[s]$ is used respectively. The simulations run for $10[s]$ what means 1000, 2000 and 4000 time steps for each case. Density and viscosity are $\rho = 1.06[g/cm3]$ and $\mu = 0.035[Poise]$.

**Initial and boundary conditions:** The initial velocity domain is $v_i \mid_{t=0} = 0$. At the inlet $\Gamma_{in}$ a parabolic constant flow profile with max velocity of $1.25$cm/s is used, given by the formula:

$$v_x \mid_{\Gamma_{in}} = -7.8125y^2 + 1.25 \tag{8}$$

The outlet is free with weakly imposed pressure equal to zero. The other two walls $\Gamma_w$ have no slip condition $v_i \mid_{\Gamma_w} = 0$.

**Analytical solution** The analytical solution should follow Eq. (8).

**Results:** The calculation of the convergence was calculated at the nodes of the coarsest mesh, to avoid any interpolation of the solution. Results for each case are sumarised in Table 1. The observed order of convergence $p$ was equal to 1.907, compatible with the theoretical order of convergence of 2 of the 2nd order backward differentiation formula (BDF) time scheme used.

##### S3.2.3 3D Womersley flow

**Problem description** : solve pulsatile Womersley flow in a 3D cylinder.

**Domain definition and space-time discretisation:** : The domain is has lenght $l = 4[cm]$ and radius $r = 0.4[cm]$. Three hexahedral meshes are used with 10179, 81432, and 651456 elements respectively. The time steps used for each case were 0.005$[s]$, 0.0025$[s]$, and 0.00125$[s]$ respectively. $10[s]$ were simulated, accounting for 2000, 4000 and 8000 time steps respectively. Density and viscosity are $\rho = 1.06[g/cm3]$ and $\mu = 0.035[Poise]$.

**Initial and boundary conditions:** The initial velocity domain is $v_i \mid_{t=0} = 0$. At the cylinder inlet a pressure boundary condition given by $P \mid_{\Gamma_{in}} = 30cos(2\pi t)[Ba]$. The outlet is free with weakly imposed pressure equal to zero. The other two walls $\Gamma_w$ have no slip condition $v_i \mid_{\Gamma_w} = 0$.

**Analytical solution:** The velocity component along the axis of the cylinder is given by:

$$v = Re\left[\frac{A}{i\omega\rho}\left(1 - \frac{J_0\left(i^{3/2}\alpha r/R\right)}{J_0(i^{3/2}\alpha)}\right)\exp(i\omega t)\right] \tag{9}$$

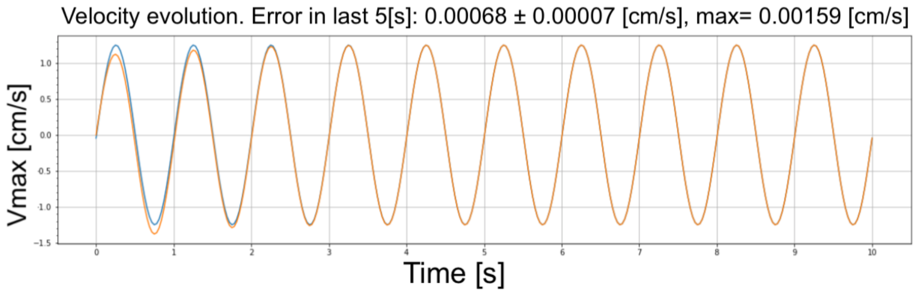


Fig. 1: Orange: Simulated maximum velocity. Blue: analytical solution.

Table 2: RMSE in $[cm/s]$ at different distances from the cylinder center at quarters of the pulsation cycle averaged radially and over the last two cycles for the finest mesh.

| Time[s] | Distance from center $[cm]$ | | | | |
|---|---|---|---|---|---|
| | 0.0 | 0.2 | 0.2 | 0.3 | 0.4 |
| x.00 | 0.000517 | 0.000313 | 0.000236 | 0.000462 | 0.0 |
| x.25 | 0.000225 | 0.000329 | 0.000311 | 0.000174 | 0.0 |
| x.50 | 0.000491 | 0.000290 | 0.000253 | 0.000471 | 0.0 |
| x.75 | 0.000244 | 0.000347 | 0.000324 | 0.000168 | 0.0 |

where:

$$A = \Delta P/L \tag{10}$$
$$\alpha = R\sqrt{\omega\rho/\mu} \tag{11}$$
$$\omega = 2\pi f \tag{12}$$

For a representation of the analytical solution refer to the blue line in Fig. 1.

**Results** : Figure 1 shows the analytical (Eq. (9)) and the simulated solution. The maximum velocity reaches expected periodic behaviour after $3[s]$. Table 2 shows the RMSE of the velocity calculation with respect to the analytical solution in the middle slice ($x = 2[cm]$) for the finest mesh. A RMSE $\epsilon = 0.0[cm/s]$ at $r = 0.4[cm]$ is explained as a Diriclhet boundary condition $v_i = 0.0[cm/s]$ is imposed there what is exactly equal to the analytical solution. The observed order of convergence is $p_{\text{obs}} = 1.82$, compatible with the theoretical order of convergence of 2 of the 2nd order BDF time scheme used. Compared to the 2D Poiseuille, a larger difference is not unexpected given the 3D Womersley is 3D and transient.

##### S3.2.4 Conclusion of the code verification tests

As the result of the code verification process we demonstrated [5]: *(a)* equations are solved correctly with at most 0.5% error with respect to the analytical solution for the Reynolds numbers representative of left ventricular assist device (LVAD) problem; *(b)* observed order of accuracy is similar to the theoretical order; *(c)* The equation coding, transformations and solution procedures are correct.

#### S3.3 Numerical calculation verification

The original model was subdivided 2 times, splitting each element into 8, to obtain meshes with $6.6M$, $53.1M$, and $425M$ elements respectively.

##### S3.3.1 Ventricular geometry with stationary boundary conditions

The goal is to estimate the stationary error and observed convergence order for the computational fluid dynamics (CFD) solver on the mesh created for the ventricular geometry. Geometry deformation, valve model, and the pump boundary condition are turned off as they depend on measurements obtained from the CFD solver. As such geometry does not have an analytical solution, the RMSE are calculated against the finest mesh computed. Results are summarised in Table 3. The observed order of convergence is $p_{\text{obs}} = 0.971$ compatible with the theoretical order of convergence of 1 provided by the first order trapezoidal time integration.

Table 3: RMSE and RMSE expressed as percentage of the maximum speed in the doamin (150$[cm/s]$) for problem specific geomety.

| halving $[-]$ | Elements $[-]$ | RMSE $[cm/s]$ | RMSE $[\%]$ | $GCI^{95\%}$ $[cm/s]$ |
|---|---|---|---|---|
| 0 | 6.6M | 1.37 | 0.91 | 1.78 |
| 1 | 53.1M | 0.511 | 0.34 | 0.66 |
| 2 | 425M | 0.0 | 0.0 | 0.0 |

##### S3.3.2 Ventricular geometry with transient boundary conditions

Here we show the RMSE for the complete model described in the main document.. As the complete model have transient boundary conditions, the RMSE is provided in time-averaged quantities. RMSEs are calculated against the finest mesh Results are summarised on Table 4. The time-averaged observed order of convergence is $\overline{p_{\text{obs}}} = 0.85$ compatible with the theoretical order of convergence of 1 provided by the first order trapezoidal time integration.

Table 4: Time-averaged RMSE ($\overline{\text{RMSE}}$), the timed-averaged RMSE% , and the time averaged $\overline{GCI^{95\%}}$ for the problem-specific gometry.

| halving $[-]$ | Elements $[-]$ | $\overline{\text{RMSE}}$ $[cm/s]$ | $\overline{\text{RMSE}}$ $[\%]$ | $\overline{GCI^{95\%}}$ $[cm/s]$ |
|---|---|---|---|---|
| 0 | 6.6M | 3.61 | 2.40 | 5.58 |
| 1 | 53.1M | 2.12 | 1.41 | 3.27 |
| 2 | 425M | 0.0 | 0.0 | 0.0 |

##### S3.3.3 Discussion of the verification results

While applicability of validation results is currently a discussion topic [6], the applicability of verification results is rarely discussed. The reason for this is probably the scarce number of verification tests that have a non-trivial analytical solution or a manufactured solution. The tests in this section are executed with Reynolds number close to the ones in the left ventricle (LV), therefore ensuring correctness of the solution procedure for an operation condition similar to the validation operation condition.

Simple, stationary physical problems as the 2D Poiseullie flow allow having small errors for a reduced computational cost. When trying to find solution to more complex transient problems (e.g. the 3D Womersley transient flow) the errors increase. The model treated in this project is not only a transient problem solved on a fine mesh, but also: *(a)* the fluid mesh is deforming, *(b)* the boundary conditions vary with the solution of the CFD solver, *(c)* it requires solving the near-ill-conditioned problem of the valve closing. Therefore, considerably larger errors were expected compared to simpler academic problem. Despite that, the numerical error remained under $2.4\%$ for the most complex use case presented.

#### References

[1] American Society of Mechanical Engineers, 2018. "Assessing Credibility of Computational Modeling through Verification and Validation: Application to Medical Devices - V V 40 - 2018". *Asme V&V 40-2018*, p. 60.

[2] Roache, P. J., 1998. *Verification and validation in computational science and engineering*, Vol. 895. Hermosa Albuquerque, NM.

[3] American Society of Mechanical Engineers, 2009. "Standard for Verification and Validation in Computational Fluid Dynamics and Heat Transfer: ASME V&V 20". *The American Society of Mechanical Engineers (ASME).*

[4] Houzeaux, G., de la Cruz, R., Owen, H., and Vázquez, M., 2013. "Parallel uniform mesh multiplication applied to a Navier-Stokes solver". *Computers and Fluids,* **80**(1), pp. 142–151.

[5] Roache, P. J., 2002. "Code verification by the method of manufactured solutions". *J. Fluids Eng.,* **124**(1), pp. 4–10.

[6] Pathmanathan, P., Gray, R. A., Romero, V. J., and Morrison, T. M., 2017. "Applicability analysis of validation evidence for biomedical computational models". *Journal of Verification, Validation and Uncertainty Quantification,* **2**(2).
